# A Quantitative Benchmark of Visual Information in Human Brain Recordings Across fMRI, MEG, and EEG

**DOI:** 10.64898/2026.01.11.698841

**Authors:** Taisei Hara, Yoshito Masai, Shinji Nishimoto

## Abstract

Noninvasive human brain recordings such as fMRI, MEG, and EEG are widely used to study visual representations, yet their relative information content has never been quantitatively benchmarked under matched experimental conditions. Here, we develop a unified decoding framework that enables fair cross-modal comparison using identical naturalistic stimuli (THINGS) and a common decoding analysis. Using a matched stimulus set, we first quantify modality performance via object-category decoding accuracy. Under these conditions, fMRI yielded the highest accuracy (mean 87%), MEG showed intermediate performance (mean 35%), and single-subject EEG achieved lower accuracy (mean 7%), while remaining reliably above chance level (1%). Notably, EEG performance improved substantially when decoding outputs were aggregated across participants in the form of a stimulus-level relational structure, yielding a group-level EEG measure (28%) with accuracy comparable to MEG. We then examined how decoding performance scales with the number of training stimuli and effective measurement time. These analyses revealed distinct efficiency–precision trade-offs across modalities: MEG and group-level EEG were most efficient for short measurement times, whereas fMRI achieved the highest asymptotic performance when sufficient data were collected. Together, these findings demonstrate that modality performance differs systematically even under tightly matched conditions and that decoding provides a principled way to assess the information content of fMRI, MEG, and EEG. The proposed unified framework offers practical guidelines for modality selection, experimental design, and data collection planning in neuroscience.

**Highlights:**

✓ Unified decoding framework enables quantitative, matched comparison of fMRI, MEG, and EEG.
✓ fMRI shows the highest decoding accuracy; MEG intermediate; EEG lower.
✓ Aggregating EEG decoding outputs into a stimulus level relational structure across subjects markedly improves performance to near MEG levels.
✓ MEG and EEG (aggregated across subjects) are most efficient for short recordings, whereas fMRI achieves the highest performance with longer measurement time.

## 1. Introduction

A central goal in systems neuroscience is to quantify how humans perceive and interpret the rich diversity of experiences encountered in daily life. Brain decoding —the process of inferring perceptual or cognitive states from neural activity— provide a powerful means of pursuing this goal (Naselaris et al., 2011). This approach has seen widely adopted in both basic research and applications such as brain-machine interfaces (Edelman et al., 2025). It has also enabled successful decoding of visual images, semantic categories, and linguistic information from human brain activity (Cichy et al., 2014; Daly, 2023; Défossez et al., 2023; Horikawa and Kamitani, 2017; Kay et al., 2008; Nishida and Nishimoto, 2018; Nishimoto et al., 2011; Santoro et al., 2017; Takagi and Nishimoto, 2023; Tang et al., 2023; Wang et al., 2024). However, despite extensive work across individual neuroimaging techniques such as fMRI, MEG, and EEG, these approaches have largely been applied in isolation (Benchetrit et al., 2023; Carlson et al., 2013; Gifford et al., 2022; Grootswagers et al., 2022; Isik et al., 2014; Nishimoto et al., 2011). As a result, it remains unclear how decodability compares across modalities under identical stimulus and analysis conditions, and no unified framework that is independent of measurement modality has been established.

Three major noninvasive neuroimaging techniques, functional magnetic resonance imaging (fMRI), magnetoencephalography (MEG), and electroencephalography (EEG), offer complementary but markedly different measurement characteristics. Because each modality captures a different physical signal, they differ substantially in spatial resolution, temporal resolution, signal-to-noise ratio (SNR), and acquisition cost. fMRI provides high-spatial resolution and relatively stable signals, but its dependence on hemodynamic responses limits temporal precision. MEG and EEG achieve millisecond-level temporal resolution but generally exhibit lower SNR compared with fMRI (Baillet, 2017; Logothetis, 2008; Schoffelen and Gross, 2009). Moreover, while fMRI and MEG require specialized facilities with high operating costs, EEG is inexpensive and well suited for data collection from large numbers of participants. These diverse characteristics play a critical role in experimental design and modality selection. However, a quantitative framework for directly comparing the representational properties of these modalities remains largely underdeveloped (Ferrante et al., 2026). Establishing such a unified benchmark is therefore essential for understanding how different neuroimaging methods capture visual information.

To address this challenge, we developed a unified analysis framework that enables direct, controlled comparison of fMRI, MEG, and EEG responses to natural images. We used the large-scale THINGS image database (Hebart et al., 2019), which contains 1,854 object categories and 22,248 images (Fig. S1a, b), together with publicly available fMRI and MEG datasets (Hebart et al., 2023) and EEG data (Grootswagers et al., 2022) collected under identical stimulus conditions (Fig. 1a). For each image, we extracted visual representations from deep neural networks (DNN), including AlexNet (Krizhevsky et al., 2012), Vision Transformer (Dosovitskiy et al., 2021) and CLIP (Radford et al., 2021). We adopted this approach given extensive evidence that hierarchical DNN features predict visual cortical responses (Cadieu et al., 2014; Eickenberg et al., 2017; Güçlü and van Gerven, 2015; Khaligh-Razavi and Kriegeskorte, 2014; Nakagi et al., 2024; Oota et al., 2022; Wen et al., 2018; Yamins et al., 2014). Using these features, we trained encoding models (Naselaris et al., 2011) to predict neural responses in each modality (Fig. 1b). Decoding was performed by identifying, for each test image, the predicted neural response that was most similar to the measured response under a Mahalanobis distance-based likelihood metric (Nishimoto et al., 2011) (Fig. 1c).

**Fig. 1.**
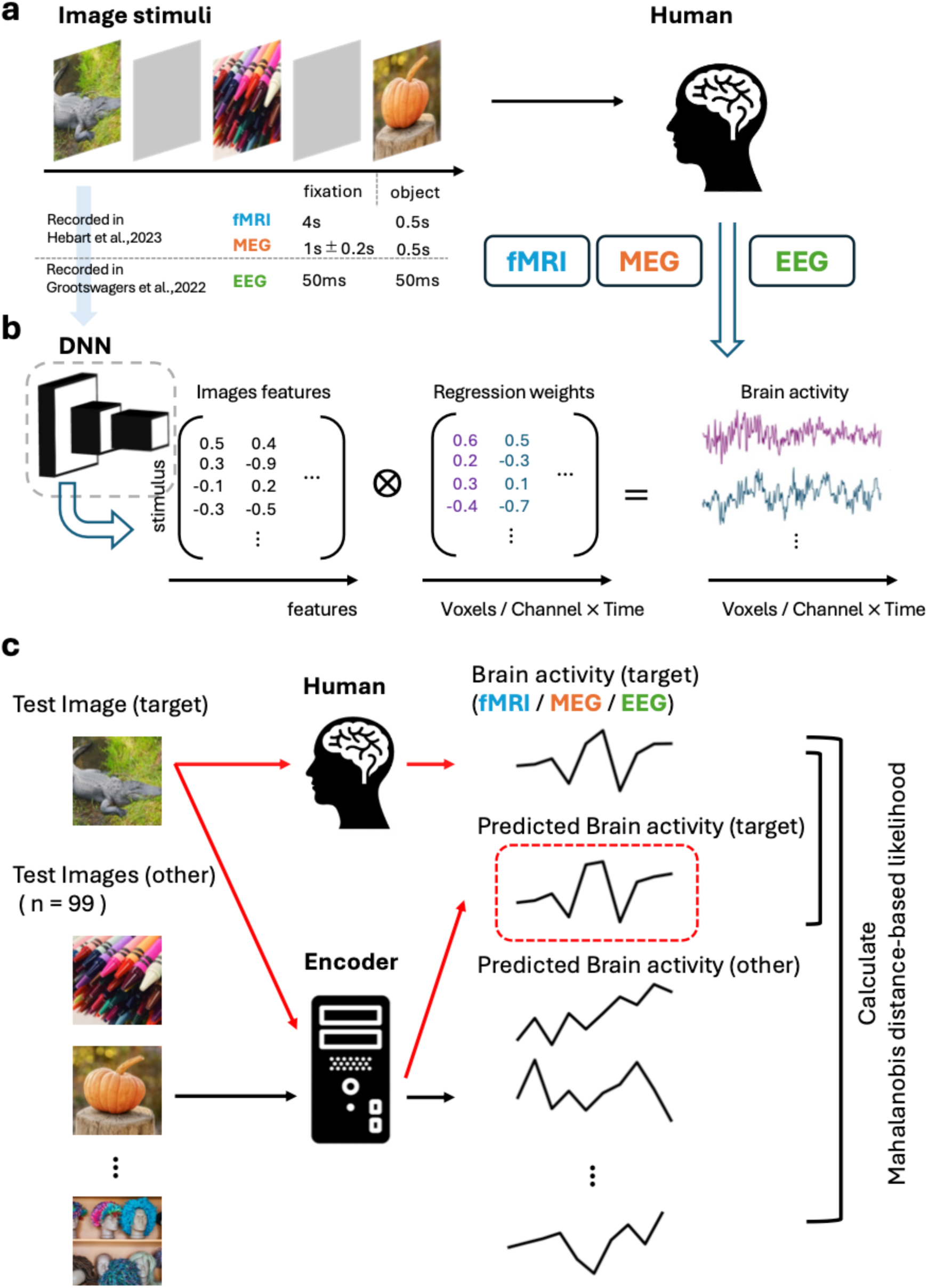
Overview of the experimental design and analysis pipeline. This figure illustrates the unified experimental and analytical framework used to compare modality performance via object-category decoding accuracy across fMRI, MEG, and EEG using an identical image stimulus set. (a) Image stimuli presentation paradigms for fMRI (0.5 s image + 4.0 s fixation), MEG (0.5 s image + 1.0 ± 0.2 s fixation), and EEG (50 ms image + 50 ms blank). (b) Encoding model framework: DNN features were extracted for each image and mapped to brain responses using ridge regression, predicting voxel-wise (fMRI) or channel × time responses (MEG/EEG). Prediction accuracy was quantified using Pearson correlation. (c) Decoding analysis using Mahalanobis distance-based likelihood: for each test image, predicted responses were compared with recorded responses, and identification accuracy was defined as the proportion of trials in which the correct image achieved the highest likelihood among 100 candidates (red dashed box). This unified analysis pipeline enables direct comparison of fMRI, MEG, and EEG under the same stimuli and a common decoding criterion. Note that the images shown here were replaced with publicly available CC0 images from THINGSplus due to copyright considerations and do not correspond to the actual experimental stimuli (Stoinski et al., 2022).

This unified framework, using identical stimuli and common encoding-decoding procedures across modalities, allowed us to systematically investigate (i) how accurately fMRI, MEG, and EEG can identify natural image categories; (ii) how decoding performance scales with the number of training stimuli and the amount of measurement time. Together, these analyses provide a quantitative basis for characterizing the visual information each modality retains and offer new insights for modality selection and experimental design in neuroscience.

## 2. Methods

### 2.1. Datasets

#### 2.1.1. Overview

All neural datasets analyzed in this study were obtained from publicly available components of the THINGS initiative (https://things-initiative.org). No new data were collected. We used multimodal neural responses (fMRI, MEG, EEG) acquired during viewing of large-scale naturalistic object images. Below we summarize the dataset specifications and preprocessing procedures performed by the data providers (Grootswagers et al., 2022; Hebart et al., 2023, 2019).

#### 2.1.2. Visual Stimuli

Visual stimuli were drawn from the THINGS image database (Hebart et al., 2019), containing 1,854 natural object categories with ≥ 12 images each (Fig. S1).

#### 2.1.3. fMRI dataset and preprocessing

We used the publicly available THINGS-fMRI dataset (Hebart et al., 2023). Functional data were acquired using a 3T Siemens Prisma system with 2-mm isotropic voxels. To stabilize head position, all sessions were acquired using individually molded headcasts. Three healthy adults (2 females, 1 male; mean age 25.33) participated.

Stimuli were presented using a fixed design: each image was shown for 0.5 s followed by 4.0 s of fixation, resulting in a stimulus onset asynchrony (SOA) of 4.5 s. To maintain attention, infrequent oddball trials consisting of artificially generated non-object images were interspersed, and participants were instructed to respond with a button-press when an oddball appeared. The training set consisted of 720 object categories with 12 images per category (8,640 images), each presented once. The test set consisted of 100 held-out images (one per category), each presented 12 times. These test images belonged to object categories present in the training set but were not included among the training images.

Standard preprocessing performed by the data providers included ICA-based denoising and regularized voxel-wise General Linear Model (GLM) estimation with optimized HRF modeling. We used the resulting single trial β estimates rather than raw BOLD time series as the fMRI response. Total measurement time per subject was 14 hours.

#### 2.1.4. MEG dataset and preprocessing

We used the publicly available THINGS-MEG dataset (Hebart et al., 2023). MEG signals were recorded using a 271-channel whole-head system at a sampling rate of 1200 Hz. To stabilize head position, individually molded headcasts were used for all sessions, and head-position tracking with marker coils was performed at the beginning and end of each run. Four healthy adults (2 females, 2 males; mean age 23.25) participated.

Stimuli were presented using a fixed design: each image was shown for 0.5 s followed by 1.0 ± 0.2 s of fixation (SOA = 1.7 ± 0.2 s). As in the fMRI dataset, occasional oddball images required a button press response to maintain attention while all other images were passively viewed. The training set consisted of 1,854 object categories with 12 images per category (22,248 images, each presented once). The test set included 200 held-out images (one per category), each presented 12 times. All test images were drawn from object categories included in the training set but were not part of the training images.

Standard preprocessing performed by the data providers included a 0.1 Hz high-pass filter, 40 Hz low-pass filter, and ICA-based artifact removal (e.g., blinks, cardiac components). Responses downsampled to 200 Hz were used for all analyses. Total measurement time per subject was 10 hours.

#### 2.1.5. EEG dataset and preprocessing

We used the publicly available THINGS-EEG dataset (Grootswagers et al., 2022). EEG data were acquired using a 64-channel BrainVision ACTiChamp system (10-10 electrode layout) at a sampling rate of 1000 Hz with Cz (vertex position in the 10-10 system) as the online reference. Fifty healthy adults participated (36 females, 14 males; mean age 20.44), but we excluded four participants because of excessive measurement noise.

Stimuli were presented using a rapid serial visual presentation (RSVP) paradigm (Grootswagers et al., 2019): each image was shown for 50 ms followed by a 50 ms blank, yielding a fixed presentation rate of 10 Hz. A central bullseye (0.5° visual angle) was continuously displayed to maintain fixation. Within each sequence, 2-5 target events (bullseye turning red for 100 ms) were inserted, and participants were instructed to press a button only for these targets while passively viewing all other images. The training set consisted of 1,854 object categories with 12 images per category (22,248 images, each presented once). The test set comprised 200 held-out images (one per category), each presented 12 times. These images came from object categories shared with the training set, while the images themselves were excluded from training.

Standard preprocessing performed by the data providers included high-pass filtering at 0.1 Hz, low-pass filtering at 100 Hz. We used responses downsampled to 250 Hz for all analyses. Total measurement time per subject was 1 hour.

### 2.2. Trial averaging

Because all modalities repeated each test set image 12 times, responses were averaged across repetitions to improve SNR (β estimates for fMRI; channel × time responses for MEG/EEG).

### 2.3. Dataset alignment and train-test splitting

Because the available images in the THINGS dataset differ slightly across modalities, we aligned all datasets to enable strict cross-modality comparison. For fMRI, we extracted responses to the 8,640 training images and 100 test images. The corresponding MEG and EEG responses were extracted for the same image sets. For MEG/EEG, we also retained the full training set of 22,248 images and 200 test images for analyses requiring larger datasets (Fig. S1a, b, 3). However, for all cross-modality comparisons, we used the overlapping set of 100 test images shared across fMRI, MEG and EEG.

### 2.4. Feature extraction with deep neural networks

We extracted image features from multiple visual models using the THINGSVision module (Muttenthaler and Hebart, 2021), including AlexNet (Krizhevsky et al., 2012), ResNet-50 (He et al., 2016), ViT-B/16 (Dosovitskiy et al., 2021), CLIP-ViT (Radford et al., 2021), DINOv2 (Oquab et al., 2023), and MambaOut (Gu et al., 2024; Huang et al., 2025). The vision-language model InternVL3.5 (Wang et al., 2025) was implemented using HuggingFace Transformers.

Images were resized, center-cropped, and normalized according to each model’s specification. For each model, we extracted features from designated layers, flattened activations, and applied PCA trained on training set images. The same PCA transform was applied to test images. All features were z-scored prior to encoding. Model details (layers, source, citations) are listed in Table S1.

### 2.5. Encoding model fitting

We constructed voxel-wise and channel-wise encoding models (Naselaris et al., 2011) to predict neural responses to visual stimuli prior to the decoding analysis. All three modalities (fMRI, MEG, EEG) were analyzed within a unified framework. For fMRI, we used voxel-wise β estimates derived from the GLM provided by the dataset authors, rather than raw BOLD time series. The analysis was restricted to visually responsive regions of interest (ROIs), specifically early and high-level visual areas such as primary visual cortex (V1-V3), fusiform face area (FFA), parahippocampal place area (PPA), and extrastriate body area (EBA), as defined by prior work (Hebart et al., 2023), and we modeled the response of each voxel separately. For MEG and EEG, we predicted channel × time responses. In both modalities, analyses were restricted to occipital sensors to focus on visually driven activity: 39 channels in the MLO/MRO/MZO groups (occipital sites) for MEG, and 17 electrodes in the P/PO/O groups (posterior occipital sites) for EEG. This design ensured that cross-modality comparisons were performed on response components that most directly reflect visual processing.

For all modalities, the feature matrix *F_E_* consisted of image features extracted from the selected layers of each DNN model. The neural response matrix *R_E_* consisted of voxel-wise β values for fMRI, or channel × time responses for MEG/EEG. Predicted responses were obtained using a linear encoding model. The predicted neural response *Ȓ_E_* was modeled by multiplying the feature matrix *F_E_* with the weight matrix *W_E_*:

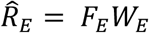

where *W_E_* is the weight matrix estimated using L2-regularized linear regression applied to the training dataset. Regularization parameters were drawn from the set 100 × 4^N^ (N = 0-9) and selected via 10-fold cross validation.

Model performance was quantified using Pearson correlation. For fMRI, voxel-wise correlations were computed and averaged across voxels. For MEG/EEG, correlations were computed at each channel and time point and then averaged across channels to obtain a time-resolved measure of encoding accuracy.

### 2.6. Decoding analysis

To quantify how much visual information each modality preserves beyond encoding accuracy, we performed an identification-based decoding analysis (Kay et al., 2008). For each test stimulus s, let r denote the observed neural response vector and *ȓ*(*s*) the response predicted from the encoding model. The likelihood of observing r given stimulus s was defined under a multivariate Gaussian noise model as (Nishimoto et al., 2011):

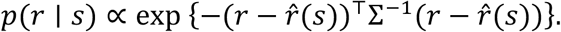

where Σ denotes the noise covariance matrix. The quadratic term in the exponent corresponds to the Mahalanobis distance between r and *ȓ*(*s*), providing a noise-normalized measure of similarity between observed and predicted responses. We estimated Σ from the residuals of the training data:

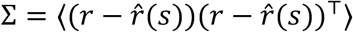

Because neural datasets are high-dimensional, Σ is frequently singular or ill-conditioned. We therefore computed Σ^-1^ using a Moore-Penrose pseudoinverse based on singular value decomposition (SVD). The regularization strength was selected via cross-validation to maximize the mean diagonal log-likelihood on hold-out validation data. Decoding was implemented as an identification procedure using similarity: for each test sample, we computed log-likelihoods for all 100 candidate images and selected the image with the highest likelihood (top-1 accuracy). Analyses were restricted to visually responsive components in each modality (fMRI: the 2,000 voxels with the highest encoding prediction accuracy on the training set within predefined visual ROIs; MEG/EEG: occipital channels in the first 200 ms after stimulus onset).

### 2.7. Model parameter optimization

We further optimized two hyperparameters of the encoding models, the number of PCA components and the DNN layer from which features were extracted, based on decoding performance. The goal of this procedure was to maximize the correspondence between DNN representations and neural responses while ensuring fair comparisons across models and stimulus counts.

For PCA dimensionality, we evaluated five settings (16, 32, 64, 128, 256). For each PCA dimension, decoding analyses were performed using features extracted from every layer of every DNN model. Identification accuracy was computed for each combination of PCA dimension, DNN layer, and stimulus subset. For each configuration, results were averaged across five random seeds, and the combination yielding the highest mean accuracy was selected as optimal. Mean accuracy was computed across all participants for fMRI and MEG, and across five randomly selected participants for EEG.

To avoid data leakage, parameter search was completely separated from evaluation (test set images, as defined in 2.3). For fMRI, 100 images from the training set were held-out exclusively for validation, and the remaining 8,540 images were used solely for model fitting. For MEG and EEG, of the 200 test set images contained in the datasets, the 100 images that were common across all modalities for cross-modality comparison were reserved for final evaluation, and only the remaining images were used during hyperparameter optimization. This design ensured strict separation between parameter search, model training, and decoding evaluation, enabling reproducible comparisons across all experimental conditions.

### 2.8. EEG aggregation strategies

To compensate for the inherently low SNR of EEG data, we evaluated five averaging and aggregation strategies across subjects. These strategies operate at different stages of the analysis pipeline and are designed to extract group-level information. Specifically, they span three processing levels: (i) averaging of response waveforms, (ii) averaging of encoding model weights, and (iii) aggregation of decoding outputs. Accordingly, they reflect increasing levels of abstraction, ranging from raw channel-level time series to stimulus-level representational structure.

In the wave-averaged EEG (wave-avg EEG) approach, we first averaged trial-averaged EEG responses for each channel across all 46 participants to obtain a single group-level response. We then used this group-level response to train a unified encoding model, thereby reducing noise arising from inter-individual differences at the level of channel time courses. In the weight-averaged EEG (weight-avg EEG) approach, we estimated encoding-weights separately for each participant and then averaged the resulting weight vectors across participants, which suppresses participant-specific noise and idiosyncratic feature-to-signal mappings. Using the encoding-weights averaged across participants, we then predicted the EEG signals of each participant and performed decoding on a per-subject basis.

We also examined subject-level averaging of decoding outputs in three different approaches. The similarity-averaged EEG (sim-avg EEG) approach computed, for each participant, a stimulus-by-stimulus similarity matrix comparing observed and predicted responses. These matrices were normalized and averaged across subjects, yielding a scale independent group-level similarity structure that preserves only the relative relationships among stimuli. In the log-likelihood averaged EEG (ll-avg EEG) approach, we averaged the log-likelihood matrices derived from the Mahalanobis distance-based decoder. This procedure relies on the fact that log-likelihoods correspond to continuous Bayesian evidence under a Gaussian noise model. This method retains more detailed information than averaging based on similarity because it aggregates continuous likelihood values before any normalization. Finally, in the rank-averaged EEG (rank-avg EEG) approach, we discretized the similarity vector into ranks for each participant. We then averaged these ranks across subjects. Because this operation depends only on ordinal information, it provides a robust, scale free aggregation that is largely insensitive to differences in absolute amplitude across individuals.

Together, these five strategies span a progression from low-level waveform averaging (wave-avg) to intermediate aggregation of encoding weights (weight-avg) and to increasingly abstract decoding-stage representations (ll-avg, sim-avg, rank-avg). Each aims to reduce subject-specific noise and trial-to-trial variability while improving the extraction of stable group-level information for EEG decoding.

### 2.9. Comparison of decoding performance under matched conditions

To evaluate modality performance under strictly controlled conditions, we conducted two complementary analyses that equated either the amount of training data or the effective measurement time across modalities. First, in the stimulus count matched analysis, we systematically manipulated the number of training images used to train the decoding models. For fMRI, the original a training set of 8,640 images (720 categories × 12 images) was subdivided into five conditions, 720, 1,440, 2,880, 5,760, and 8,640 images, corresponding to 1, 2, 4, 8, or 12 images per category. For each of the four subset conditions, we generated five independent training subsets by randomly sampling images within each category using different random seeds. The same subsampling procedure was applied to the MEG and EEG datasets, where a training set of 22,248 images (1,854 categories × 12 images) was treated as an additional data condition. All reported accuracies represent the average across the five independently generated subsets for each sample size.

In the measurement time matched analysis, we accounted for the substantial differences in acquisition duration across fMRI, MEG, and EEG by comparing decoding performance as a function of equivalent measurement time. For each modality, we computed the total measurement time per participant (approximately 840 min for fMRI, 600 min for MEG, and 60 min for EEG) and converted these durations into the corresponding numbers of training set images based on the measurement time per image. Here, measurement time per image was defined as the total measurement time per participant divided by the total number of stimulus images presented per participant for that modality. Using this conversion, we constructed matched time budget conditions for each modality. For fMRI, we evaluated seven training conditions matched for measurement time (180, 360, 720, 1,440, 2,880, 5,760, and 8,640 images). The 180 and 360 image conditions were generated by first selecting one representative image per category to form a pool of 720 images and then randomly sampling 180 or 360 images from this pool; five independent subsets were again created for each condition. For MEG, the smallest condition (180 images) was excluded due to insufficient trials, and for EEG the 360 images condition was excluded; both MEG and EEG modalities additionally included the condition with 22,248 images. As in the stimulus count analysis, five random seeds were used to generate independent training subsets for each condition. Because EEG provided only 60 minutes of data per participant, the later portion of the measurement time curve could not be observed directly. We therefore fitted a saturating exponential function, *y* = *A*(1 − *e*^−*Bx*^) + *C*, to the empirical EEG points to extrapolate its asymptotic trend.

## 3. Results

### 3.1. Encoding model construction across fMRI, MEG, and EEG

To quantitatively compare the representational characteristics of the three measurement modalities, we first constructed the unified analysis pipeline illustrated in Fig. 1. Using an identical stimulus set from the THINGS database (1,854 natural object categories; Fig. S1a, b), we applied common encoding and Mahalanobis distance-based likelihood decoding analyses to fMRI, MEG, and EEG (Fig. 1b, c). For each modality, model parameters were optimized via cross-validation, and encoding models were trained using feature representations extracted from each layer of DNN (see Supplementary Materials). Analyses were first conducted using AlexNet (Krizhevsky et al., 2012) as a reference model.

### 3.2. Modality differences in decoding performance

Using the encoding models constructed above, we next performed decoding analyses to quantitatively compare identification performance across the three modalities (fMRI, MEG, EEG). Under the stimulus count matched condition (N = 8,640 training images), fMRI achieved the highest decoding accuracy, demonstrating a clear advantage for object-category identification (Kay et al., 2008). Specifically, fMRI reached an accuracy of 87.3 ± 8.3%, compared with 34.5 ± 24.0% for MEG and 6.8 ± 8.0% for single-subject EEG (Fig. 2e). Examination of representational similarity matrices further illustrated these differences: fMRI exhibited a strong diagonal structure with consistently high-likelihoods for correct categories, reflecting stable and reliable decoding performance (Fig. 2a). In contrast, MEG and EEG showed greater variability in their diagonal components, with a larger proportion of errors in nondiagonal elements (Fig. 2b, c). Although single-subject EEG accuracy was generally low, 34 of the 46 participants exceeded the 1% chance level. This indicates that the EEG signal still contains weak but detectable stimulus specific information. Overall, the decoding performance followed a clear hierarchy of fMRI > MEG > EEG, mirroring the relative information content available in each modality.

**Fig. 2.**
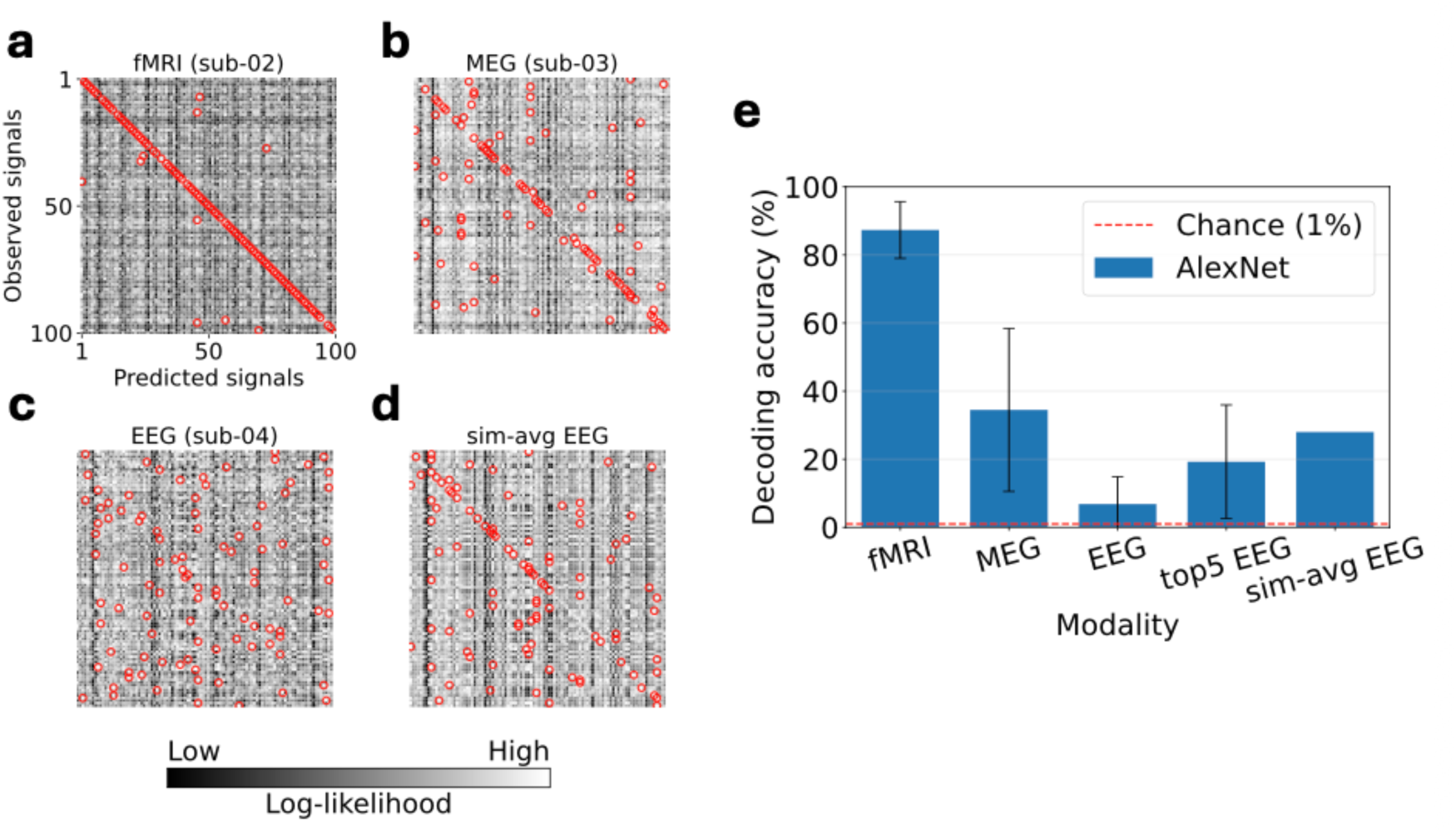
Decoding performance and similarity structure across neuroimaging modalities. (a-d) Representational similarity matrices for fMRI (a), MEG (b), EEG (c), and group-level EEG (sim-avg EEG, d). Higher values along the diagonal indicate stronger consistency between the predicted and observed responses for the target images; corresponding decoding accuracies were fMRI = 90%, MEG = 44%, EEG = 8%, and sim-avg EEG = 28%. Red circles indicate the highest likelihood. (e) Comparison of decoding performance across fMRI, MEG, EEG, and group-level EEG (sim-avg EEG).

### 3.3. EEG redundancy and group-level averaging

In single-subject EEG, decoding accuracy remained low, prompting us to examine whether weak but redundant information might nevertheless be present in the likelihood distribution. As an auxiliary measure, we computed top-5 accuracy. Whereas top-1 accuracy reached only 6.8%, top-5 accuracy increased to 19.3% ± 16.7%, indicating that individual EEG responses contain diffuse, yet interpretable high-likelihood evidence distributed across multiple candidate stimuli. We then asked whether such redundant evidence could be made more informative at the group-level. Averaging the response similarity matrices across subjects (sim-avg EEG) led to a substantial improvement in decoding performance (Fig. 2d, e). When we varied the number of participants, accuracy increased monotonically with sample size (Fig. S7). This pattern indicates that variability across individuals is a major noise source and that group-level aggregation effectively extracts stable structure from EEG responses. We also provide a systematic comparison of several group-level aggregation strategies (See Supplementary Materials).

### 3.4. Decoding performance as a function of training dataset size

To compare modality performance under strictly matched conditions, we evaluated decoding accuracy while varying the number of training images used for encoding model construction. For the fMRI dataset, the full set of 8,640 training images was subdivided into multiple stimulus count conditions, and the same subdivision rule was applied to the MEG and EEG datasets. In addition, for MEG and EEG, we trained the encoding models using the dataset of 22,248 images. All models were trained using common procedures described above, and optimal parameters were determined by cross-validation within each stimulus count condition.

Figure 3a summarizes how decoding accuracy changed as a function of the number of training images for each modality. For all modalities, performance increased as the number of training stimuli increased, with fMRI showing the steepest improvement. When only 720 training images were used, modality differences were modest, but as the dataset size increased the separation between modalities became increasingly pronounced. MEG and EEG also showed improved performance with larger training sets, and MEG continued to benefit, although the improvement was more gradual, when trained on the full set of 22,248 images. For example, MEG achieved a mean accuracy of 53.0% with 22,248 images. By comparison, its accuracy was comparable to that of fMRI trained on only 1,440 images (47.1%). In contrast, single-subject EEG decoding accuracy remained substantially lower than the other modalities and showed only limited gains even with increased stimulus counts, reaching an average of 8.8% at 22,248 images.

**Fig. 3.**
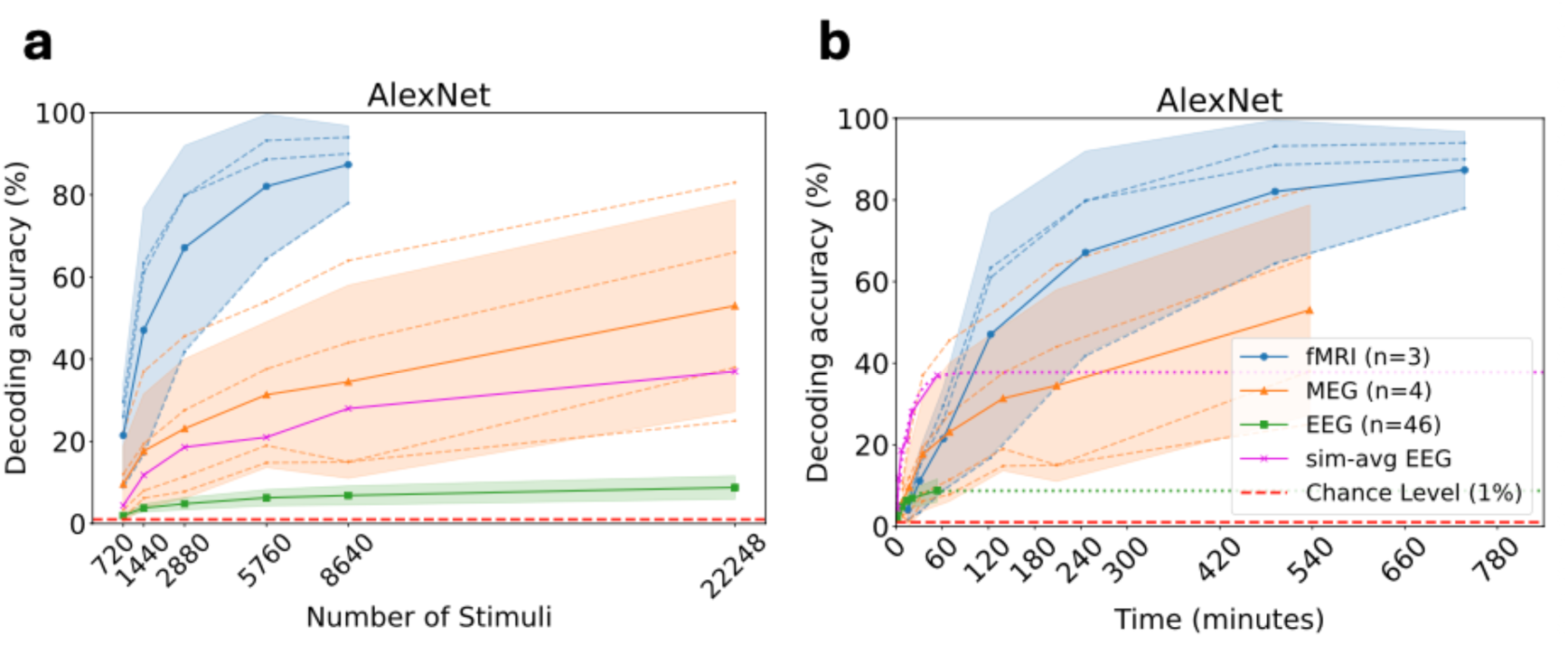
Decoding performance as a function of training sample size and effective measurement time across modalities. (a) Decoding accuracy increased monotonically with the number of training images for all modalities. fMRI exhibited the steepest improvement and reached near saturation with fewer stimuli, whereas single-subject EEG showed only modest gains. Dashed lines indicate individual participants (fMRI, MEG), solid lines represent participant averages, shaded areas indicate 95% confidence intervals (CI), and the red dashed line marks chance level (1%). (b) Decoding accuracy plotted as a function of effective measurement time. MEG and group-level EEG were most efficient for short acquisition durations, whereas fMRI achieved the highest asymptotic performance when sufficient measurement time was available. In addition, because the EEG training set was collected over a much shorter measurement duration than the other modalities, the later portion of the curve was supplemented by fitting a saturating function to the available data. Shaded areas indicate 95% CI, and red dashed lines denote chance level.

However, when stimulus-level similarity structures for each subject were averaged (sim-avg EEG), EEG achieved MEG-like performance regardless of stimulus count, reaching 37% at 22,248 images, slightly exceeding the MEG accuracy at 8,640 images (34.5%). This result strongly supports the effectiveness of aggregation based on similarity for extracting stable information from EEG data. It should also be noted that fMRI and MEG exhibited wider 95% confidence intervals due to their smaller numbers of participants (3 for fMRI, 4 for MEG), leading to greater variability in estimated decoding accuracy.

### 3.5. Decoding performance as a function of effective measurement time

To compare decoding performance across modalities on a common temporal scale, we next evaluated accuracy as a function of effective measurement time. Because fMRI, MEG, and EEG differ substantially in the amount of measurement time required per stimulus, we converted each training dataset into normalized units of measurement time. Specifically, for each modality we computed the number of training images obtainable given its total recording duration per participant (fMRI: 840 min; MEG: 600 min; EEG: 60 min), considering both the number of available trials and the measurement time per image. This procedure enabled a unified assessment of decoding performance per unit of measurement time while preserving modality specific experimental constraints.

Figure 3b summarizes the results. Although fMRI requires considerably longer measurement times, its decoding accuracy increased sharply with additional measurement time. MEG, by contrast, achieved performance comparable to or exceeding that of fMRI under short duration conditions. Within the first 60 minutes of data collection, MEG outperformed fMRI. EEG decoding accuracy was generally lower than the other modalities in the single-subject condition; however, sim-avg EEG consistently exceeded both fMRI and MEG under conditions equivalent to 60 minutes of measurement time. For example, whereas MEG achieved 34.5% accuracy at the ∼210 minutes (≈8,640 images) condition, sim-avg EEG reached 37% accuracy using only 60 minutes of EEG data, highlighting the marked efficiency gain obtained through group-level averaging. At longer durations, fMRI showed a clear advantage: around 180 minutes, fMRI surpassed all MEG and group-level EEG conditions and ultimately reached accuracy levels near 90% with extended measurement time. Due to the small number of participants in the fMRI (N = 3) and MEG (N = 4) datasets, the corresponding 95% confidence intervals were relatively wide, reflecting greater variability across participants.

To further characterize how modality performance varies as a function of measurement time, we summarized the “winner” at several interpretable time thresholds (Fig. 3b). In very short sessions (< 20 min), sim-avg EEG was the strongest modality, outperforming MEG and remaining comparable to fMRI despite the short acquisition duration. In the 20-60 min range, MEG began to surpass fMRI, while sim-avg EEG continued to provide the highest performance among all modalities. In the 60-110 min range, fMRI overtook MEG, yet sim-avg EEG still yielded the strongest performance overall. At approximately 180 min, fMRI became the single best performing modality, exceeding both MEG and group-level EEG. Finally, at very long durations (≈300 min), MEG ultimately surpassed sim-avg EEG, although fMRI maintained the highest performance across all modalities. Taken together, these comparisons across time thresholds reveal a dynamic performance hierarchy in which sim-avg EEG dominates in very short sessions, MEG becomes competitive at intermediate durations, and fMRI achieves the strongest performance with extended measurement times.

### 3.6. Architecture generalization of encoding and decoding patterns

To determine whether the encoding and decoding patterns observed with AlexNet generalize beyond a single architecture, we next evaluated a broader set of DNN that differ in structure, learning paradigm, and representational properties. These included a deep residual convolutional neural network (CNN), ResNet-50 (He et al., 2016), a Vision Transformer with self-attention, ViT (Dosovitskiy et al., 2021), a state space sequence model optimized for temporal integration, MambaOut (Gu et al., 2024; Huang et al., 2025), a vision-language model trained with contrastive objectives, CLIP (Radford et al., 2021), a self-supervised model, DINOv2 (Oquab et al., 2023), and a large-scale vision-language model, InternVL3.5 (Wang et al., 2025). As part of this analysis, we examined whether encoding patterns generalize across architectures (see Supplementary Materials).

In decoding analyses, fMRI consistently achieved the highest identification accuracy across all models, with MEG following and single-subject EEG performing lowest (Fig. 4). Notably, sim-avg EEG closely approached MEG-level performance for every model. This result indicates that EEG contains reliable stimulus information that becomes evident when aggregated across participants. The vision-language model InternVL3.5 showed lower decoding performance across all modalities compared with the other models. Importantly, the relative modality hierarchy observed with AlexNet, fMRI > MEG ≈ sim-avg EEG > single-subject EEG, was preserved across all models and remained stable under variations in stimulus count and effective measurement time (Fig. S5, S6). In early measurement time regimes, both MEG and sim-avg EEG outperformed fMRI, whereas after approximately 180 minutes, fMRI surpassed all other conditions and ultimately reached the highest performance overall. Together, these findings indicate that the observed modality differences do not arise from idiosyncrasies of specific network architectures or training paradigms but instead reflect intrinsic measurement characteristics of each modality. The fact that these cross-modality performance patterns were replicated across a wide range of models underscores the robustness and generality of the proposed modality comparison framework.

**Fig. 4.**
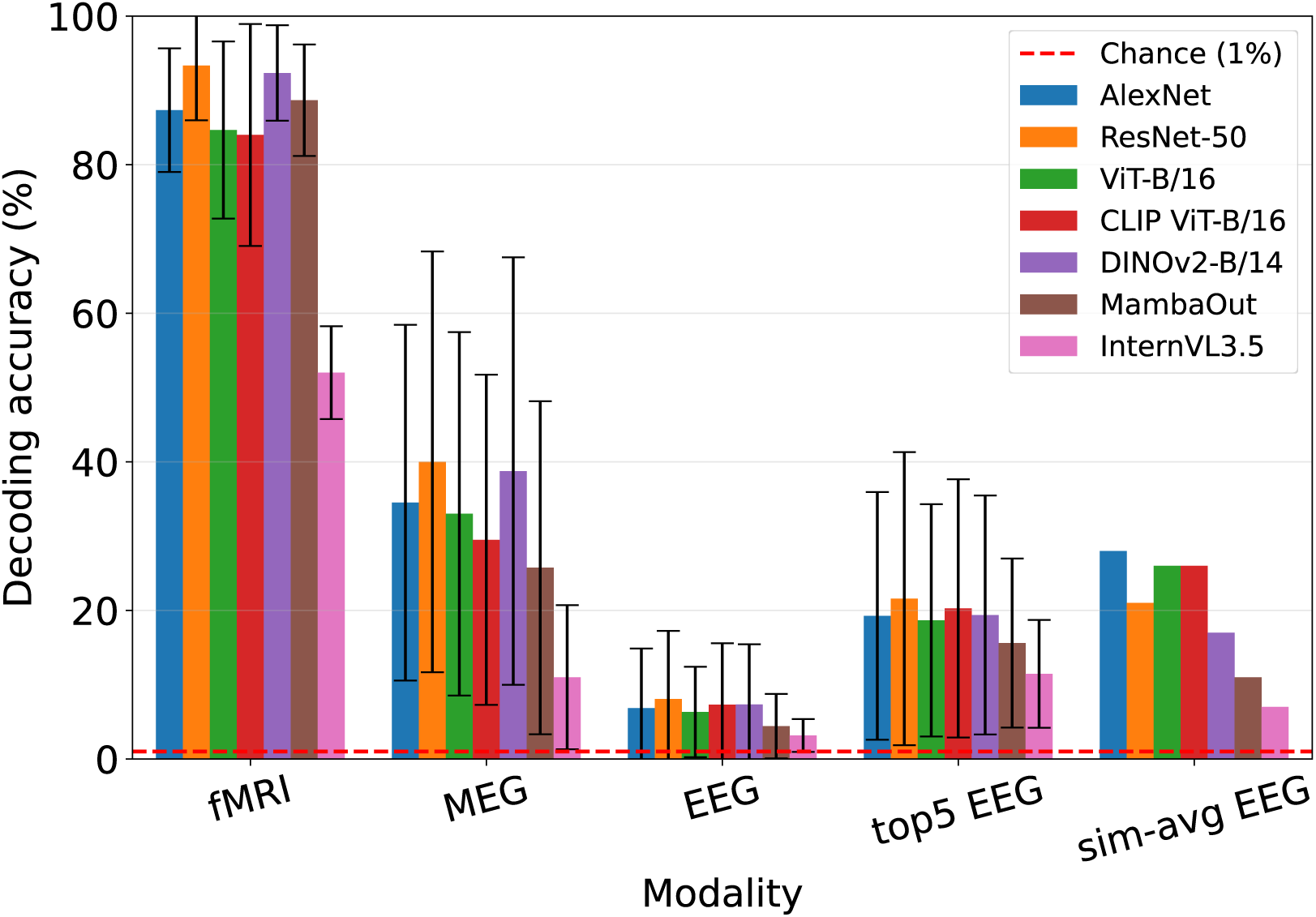
Comparison of decoding performance across fMRI, MEG, EEG, and group-level EEG. This figure summarizes the decoding accuracy for all modalities across multiple visual processing models (CNNs, Transformers, self-supervised models, and vision-language models). Bars represent the mean ± SD across participants, with model-specific colors defined in the legend. The primary metric is top-1 accuracy. For EEG, top-5 accuracy is additionally shown as an auxiliary measure of redundant decoding evidence. fMRI showed the highest performance, followed by MEG, sim-avg EEG, top-5 EEG, and single-subject EEG.

## 4. Discussion

### 4.1. Resource dependent trade-offs in decodability across fMRI, MEG, and EEG

In this study, we established a unified analysis framework that enables direct comparisons of fMRI, MEG, and EEG representations under tightly controlled conditions. This framework uses an identical stimulus set and a common encoding and decoding pipeline. Although many prior decoding studies have demonstrated strong performance within a single modality (Horikawa and Kamitani, 2017; Nishimoto et al., 2011; Takagi and Nishimoto, 2023; Tang et al., 2023), systematic cross-modality comparisons have remained limited, largely because datasets in which fMRI, MEG, and EEG are collected using matched stimulus sets are scarce. Our framework addresses this gap by providing a principled, fair comparison in which decoding accuracy is used as an operational measure of the stimulus related information available in each modality. We found a clear trade off across fMRI, MEG, and EEG between measurement resources and reliably extractable stimulus related information. Here, measurement resources were defined by acquisition time per participant and sample size. Accordingly, rather than considering any single modality as universally optimal, the relative utility of each modality depends strongly on constraints imposed by experimental design and data acquisition.

In applied domains such as brain-machine interfaces and neuromarketing, practical constraints including available recording time, the number of participants that can be recruited, and cost often effectively determine the attainable amount of information. When sufficient acquisition time can be allocated to each participant, fMRI can ultimately reach the highest asymptotic performance, making it advantageous for applications that prioritize representational fidelity and information richness. In contrast, when rapid access to stimulus-related information is required within limited measurement time, MEG represents a more time-efficient option. Furthermore, although EEG provides only limited extractable information at the single-subject level, it can become competitive when data are collected from many participants. In such cases, aggregating evidence across participants as a stimulus-level relational structure (sim-avg EEG) enables reliable extraction of stable stimulus-related structure. In addition, EEG is relatively inexpensive and portable compared with fMRI and MEG, making it particularly useful for large-scale data collection and deployment in real-world or resource-limited settings.

### 4.2. Noise dependent optimality of group-level aggregation strategies

The group-level analyses revealed a striking dissociation in the optimal aggregation strategy across modalities (Figs. 3, S7-S10), suggesting that the noise structure inherent to each measurement technique determines the representational abstraction level at which information can be reliably pooled. In EEG, where single-subject responses exhibit low SNR, large amplitude variability, and substantial difference across individuals, averaging at the level of raw waveforms or encoding weights failed to produce meaningful improvements. Moreover, likelihood-based averaging eventually degraded with increasing the number of subjects, reflecting the instability of continuous valued evidence when likelihood scales differ across individuals. This instability is further exacerbated by the fact that participants differ not only in the overall scale of likelihood values but also in their noise magnitudes, causing high-noise subjects to disproportionately influence the aggregated likelihoods. In contrast, averaging based on similarity and rank consistently provided robust improvements (Fig. S7-S9), indicating that these methods generate representations based solely on relational structure and are therefore less sensitive to amplitude scaling and subject specific variability. As a result, the stable relational structure preserved in EEG becomes more clearly expressed.

MEG showed a different pattern. The intermediate SNR and reduced differences in scale across participants allowed log-likelihood averaging to retain more detailed information, outperforming methods based on similarity and rank. This suggests that MEG contains sufficiently stable continuous evidence such that discarding absolute likelihood values is suboptimal. fMRI, with the highest SNR and most stable response scaling, showed comparable improvements across averaging strategies, indicating that its representational structure is robust to the level of abstraction at which aggregation occurs (Fig. S10). Together, these findings support the idea that optimal aggregation depends on the modality’s noise structure, in which (i) modalities with unstable amplitude scales (EEG) benefit from scale invariant relational averaging, (ii) modalities with moderate-SNR but stable likelihood structure (MEG) benefit from continuous evidence aggregation, and (iii) high-SNR modalities (fMRI) remain robust across aggregation levels. However, note that, unlike EEG which included 46 participants, the fMRI and MEG results were obtained from much smaller samples (3 and 4 participants, respectively).

This principle unifies the observed modality differences and highlights that the optimal group-level strategy is not universal but tightly coupled to the modality specific noise landscape. More broadly, these results suggest that group-level decoding in low-SNR modalities should prioritize relational representations (similarity/rank), whereas medium-SNR modalities should leverage continuous Bayesian evidence (log-likelihood). These noise aware design principles may extend to brain activity data sets beyond vision. They may also be applicable to other low or medium SNR modalities, such as fNIRS (Klein, 2024; Pereira et al., 2023; Pinti et al., 2020).

### 4.3. Limitations

Although our framework provides a unified comparison across modalities, several practical constraints should be acknowledged. First, the number of participants differed substantially across modalities (fMRI: 3; MEG: 4; EEG: 46), which inevitably leads to wider confidence intervals and less stable decodability estimates in fMRI and MEG. Second, it remains unclear whether the measurement procedures were fully appropriate for each modality. In particular, the EEG data were collected using an RSVP paradigm with very brief stimulus exposures. Such rapid presentation may suppress or truncate early visual responses through forward and backward masking, thereby reducing recoverable low-level information and contributing to lower performance (Grootswagers et al., 2019). Third, our comparisons focused exclusively on visual modalities. Other sensory systems such as audition, olfaction, or language processing were not examined. These modalities differ substantially in temporal dynamics, representational structure, and sensorimotor coupling. Extending the framework to such domains will be essential for understanding how modality specific constraints shape decodability.

## Conclusion

In this study, we developed a unified framework that enables direct comparison of visual representations across fMRI, MEG, and EEG using identical stimuli and a common encoding and decoding pipeline (Fig. 1). Decoding analyses revealed a consistent performance hierarchy of fMRI > MEG ≈ sim-avg EEG > single-subject EEG, under matched stimulus sets (Fig. 2, 4). Analyses of stimulus count and measurement time further showed that MEG and group-level EEG are most efficient in regimes with short measurement durations, whereas fMRI achieves the highest accuracy with extended measurement time (Fig. 3). Importantly, the group-level EEG results demonstrate that scalable EEG measurements with relatively low-cost can yield MEG-like performance through appropriate aggregation. This provides a practical alternative when fMRI or MEG are limited by cost or accessibility. These findings broaden the range of viable experimental designs by clarifying how measurement precision, scalability, and aggregation strategies jointly shape decodability. Taken together, our results provide quantitative guidance for modality selection and experiment planning. Future work may extend this framework to additional stimulus domains and task paradigms.

## Acknowledgements

This study was supported by KAKENHI JP24H00619 and JST JPMJCR24U2.

## Reference

Baillet, S., 2017. Magnetoencephalography for brain electrophysiology and imaging. Nat. Neurosci. 20, 327–339.

Benchetrit, Y., Banville, H., King, J.-R., 2023. Brain decoding: toward real-time reconstruction of visual perception. arXiv [eess.IV].

Cadieu, C.F., Hong, H., Yamins, D.L.K., Pinto, N., Ardila, D., Solomon, E.A., Majaj, N.J., DiCarlo, J.J., 2014. Deep neural networks rival the representation of primate IT cortex for core visual object recognition. PLoS Comput. Biol. 10, e1003963.

Carlson, T., Tovar, D.A., Alink, A., Kriegeskorte, N., 2013. Representational dynamics of object vision: the first 1000 ms. J. Vis. 13, 1–19.

Cichy, R.M., Pantazis, D., Oliva, A., 2014. Resolving human object recognition in space and time. Nat. Neurosci. 17, 455–462.

Daly, I., 2023. Neural decoding of music from the EEG. Sci. Rep. 13, 624.

Défossez, A., Caucheteux, C., Rapin, J., Kabeli, O., King, J.-R., 2023. Decoding speech perception from non-invasive brain recordings. Nat. Mach. Intell. 5, 1097–1107.

Dosovitskiy, A., Beyer, L., Kolesnikov, A., Weissenborn, D., Zhai, X., Unterthiner, T., Dehghani, M., Minderer, M., Heigold, G., Gelly, S., Uszkoreit, J., Houlsby, N., 2021. An Image is Worth 16x16 Words: Transformers for Image Recognition at Scale, in: International Conference on Learning Representations (ICLR).

Edelman, B.J., Zhang, S., Schalk, G., Brunner, P., Muller-Putz, G., Guan, C., He, B., 2025. Non-invasive brain-computer interfaces: State of the art and trends. IEEE Rev. Biomed. Eng. 18, 26–49.

Eickenberg, M., Gramfort, A., Varoquaux, G., Thirion, B., 2017. Seeing it all: Convolutional network layers map the function of the human visual system. Neuroimage 152, 184–194.

Ferrante, M., Boccato, T., Rashkov, G., Toschi, N., 2026. Towards neural foundation models for vision: Aligning EEG, MEG, and fMRI representations for decoding, encoding, and modality conversion. Inf. Fusion 126, 103650.

Gifford, A.T., Dwivedi, K., Roig, G., Cichy, R.M., 2022. A large and rich EEG dataset for modeling human visual object recognition. Neuroimage 264, 119754.

Grootswagers, T., Robinson, A.K., Carlson, T.A., 2019. The representational dynamics of visual objects in rapid serial visual processing streams. Neuroimage 188, 668–679.

Grootswagers, T., Zhou, I., Robinson, A.K., Hebart, M.N., Carlson, T.A., 2022. Human EEG recordings for 1,854 concepts presented in rapid serial visual presentation streams. Sci. Data 9, 3.

Gu, A., Dao, T., Others, 2024. Mamba: Linear-Time Sequence Modeling with Selective State Spaces. Presented at the Proceedings of the Conference on Language Modeling (COLM).

Güçlü, U., van Gerven, M.A.J., 2015. Deep neural networks reveal a gradient in the complexity of neural representations across the ventral stream. J. Neurosci. 35, 10005–10014.

He, K., Zhang, X., Ren, S., Sun, J., 2016. Deep Residual Learning for Image Recognition, in: Proceedings of the IEEE Conference on Computer Vision and Pattern Recognition (CVPR). pp. 770–778.

Hebart, M.N., Contier, O., Teichmann, L., Rockter, A.H., Zheng, C.Y., Kidder, A., Corriveau, A., Vaziri-Pashkam, M., Baker, C.I., 2023. THINGS-data, a multimodal collection of large-scale datasets for investigating object representations in human brain and behavior. Elife 12. 10.7554/eLife.82580

Hebart, M.N., Dickter, A.H., Kidder, A., Kwok, W.Y., Corriveau, A., Van Wicklin, C., Baker, C.I., 2019. THINGS: A database of 1,854 object concepts and more than 26,000 naturalistic object images. PLoS One 14, e0223792.

Horikawa, T., Kamitani, Y., 2017. Generic decoding of seen and imagined objects using hierarchical visual features. Nat. Commun. 8, 15037.

Huang, Z., Zhai, X., Chen, X., Xu, Y., Pang, R., Le, Q.V., Wang, Z., 2025. MambaOut: Layer Scale-free Mamba, in: ArXiv Preprint ArXiv:2405. 16673. Presented at the Proceedings of the IEEE/CVF Conference on Computer Vision and Pattern Recognition (CVPR).

Isik, L., Meyers, E.M., Leibo, J.Z., Poggio, T., 2014. The dynamics of invariant object recognition in the human visual system. J. Neurophysiol. 111, 91–102.

Kay, K.N., Naselaris, T., Prenger, R.J., Gallant, J.L., 2008. Identifying natural images from human brain activity. Nature 452, 352–355.

Khaligh-Razavi, S.-M., Kriegeskorte, N., 2014. Deep supervised, but not unsupervised, models may explain IT cortical representation. PLoS Comput. Biol. 10, e1003915.

Klein, F., 2024. Optimizing spatial specificity and signal quality in fNIRS: an overview of potential challenges and possible options for improving the reliability of real-time applications. Front. Neuroergonomics 5, 1286586.

Krizhevsky, A., Sutskever, I., Hinton, G.E., 2012. ImageNet Classification with Deep Convolutional Neural Networks, in: Advances in Neural Information Processing Systems 25 (NIPS 2012). pp. 1106–1114.

Logothetis, N.K., 2008. What we can do and what we cannot do with fMRI. Nature 453, 869–878.

Muttenthaler, L., Hebart, M.N., 2021. THINGSvision: A Python toolbox for streamlining the extraction of activations from deep neural networks. Front. Neuroinform. 15, 679838.

Nakagi, Y., Matsuyama, T., Koide-Majima, N., Yamaguchi, H.Q., Kubo, R., Nishimoto, S., Takagi, Y., 2024. Unveiling multi-level and multi-modal semantic representations in the human brain using large language models. Presented at the Proceedings of the 2024 Conference on Empirical Methods in Natural Language Processing, Association for Computational Linguistics, pp. 20313–20338.

Naselaris, T., Kay, K.N., Nishimoto, S., Gallant, J.L., 2011. Encoding and decoding in fMRI. Neuroimage 56, 400–410.

Nishida, S., Nishimoto, S., 2018. Decoding naturalistic experiences from human brain activity via distributed representations of words. Neuroimage 180, 232–242.

Nishimoto, S., Vu, A.T., Naselaris, T., Benjamini, Y., Yu, B., Gallant, J.L., 2011. Reconstructing visual experiences from brain activity evoked by natural movies. Curr. Biol. 21, 1641–1646.

Oota, S.R., Arora, J., Rowtula, V., Gupta, M., Bapi, R.S., 2022. Visio-Linguistic Brain Encoding, in: Proceedings of the 29th International Conference on Computational Linguistics. pp. 116–133.

Oquab, M., Darcet, T., Moutakanni, T., Valko, M., Bojanowski, P., Luc, P., Joulin, Alan, Joulin, Armand, Misra, I., Jegou, H., Mairal, J., 2023. DINOv2: Learning Robust Visual Features without Supervision. arXiv preprint arXiv:2304. 07193.

Pereira, J., Direito, B., Lührs, M., Castelo-Branco, M., Sousa, T., 2023. Multimodal assessment of the spatial correspondence between fNIRS and fMRI hemodynamic responses in motor tasks. Sci. Rep. 13, 2244.

Pinti, P., Tachtsidis, I., Hamilton, A., Hirsch, J., Aichelburg, C., Gilbert, S., Burgess, P.W., 2020. The present and future use of functional near-infrared spectroscopy (fNIRS) for cognitive neuroscience. Ann. N. Y. Acad. Sci. 1464, 5–29.

Radford, A., Kim, J.W., Hallacy, C., Ramesh, A., Goh, G., Agarwal, S., Sastry, G., Askell, A., Mishkin, P., Clark, J., Krueger, G., Sutskever, I., 2021. Learning Transferable Visual Models From Natural Language Supervision, in: The 43rd Annual Conference of the Cognitive Science Society (CogSci 2021).

Santoro, R., Moerel, M., De Martino, F., Valente, G., Ugurbil, K., Yacoub, E., Formisano, E., 2017. Reconstructing the spectrotemporal modulations of real-life sounds from fMRI response patterns. Proc. Natl. Acad. Sci. U. S. A. 114, 4799–4804.

Schoffelen, J.-M., Gross, J., 2009. Source connectivity analysis with MEG and EEG. Hum. Brain Mapp. 30, 1857–1865.

Stoinski, L.M., Perkuhn, J., Hebart, M.N., 2022. THINGSplus: New norms and metadata for the THINGS database of 1,854 object concepts and 26,107 natural object images. PsyArXiv. 10.31234/osf.io/exu9f

Takagi, Y., Nishimoto, S., 2023. High-Resolution Image Reconstruction with Latent Diffusion Models From Human Brain Activity, in: Proceedings of the IEEE/CVF Conference on Computer Vision and Pattern Recognition (CVPR). IEEE / Computer Vision Foundation, pp. 14453–14463.

Tang, J., LeBel, A., Jain, S., Huth, A.G., 2023. Semantic reconstruction of continuous language from non-invasive brain recordings. Nat. Neurosci. 26, 858–866.

Wang, B., Xu, X., Zhang, L., Xiao, B., Wu, X., Chen, J., 2024. Semantic reconstruction of continuous language from meg signals, in: ICASSP 2024 - 2024 IEEE International Conference on Acoustics, Speech and Signal Processing (ICASSP). Presented at the ICASSP 2024 - 2024 IEEE International Conference on Acoustics, Speech and Signal Processing (ICASSP), IEEE, pp. 2190–2194.

Wang, Weiyun, Gao, Z., Gu, L., Pu, H., Cui, L., Wei, X., Liu, Z., Jing, L., Ye, S., Shao, J., Wang, Z., Chen, Z., Zhang, H., Yang, G., Wang, H., Wei, Q., Yin, J., Li, W., Cui, E., Chen, G., Ding, Z., Tian, C., Wu, Z., Xie, J., Li, Z., Yang, B., Duan, Y., Wang, X., Hou, Z., Hao, H., Zhang, T., Li, S., Zhao, X., Duan, H., Deng, N., Fu, B., He, Y., Wang, Y., He, C., Shi, B., He, J., Xiong, Y., Lv, Han, Wu, L., Shao, W., Zhang, K., Deng, H., Qi, B., Ge, J., Guo, Q., Zhang, W., Zhang, S., Cao, M., Lin, J., Tang, K., Gao, J., Huang, H., Gu, Y., Lyu, C., Tang, H., Wang, R., Lv, Haijun, Ouyang, W., Wang, L., Dou, M., Zhu, X., Lu, T., Lin, D., Dai, J., Su, W., Zhou, B., Chen, K., Qiao, Y., Wang, Wenhai, Luo, G., 2025. InternVL3.5: Advancing open-source multimodal models in versatility, reasoning, and efficiency. arXiv [cs.CV].

Wen, H., Shi, J., Chen, W., Liu, Z., 2018. Deep residual network predicts cortical representation and organization of visual features for rapid categorization. Sci. Rep. 8, 3752.

Yamins, D.L.K., Hong, H., Cadieu, C.F., Solomon, E.A., Seibert, D., DiCarlo, J.J., 2014. Performance-optimized hierarchical models predict neural responses in higher visual cortex. Proc. Natl. Acad. Sci. U. S. A. 111, 8619–8624.

