## Supplementary materials for "A Quantitative Benchmark of Visual Information in Human Brain Recordings Across fMRI, MEG, and EEG"

### Supplementally results

#### Encoding model results across fMRI, MEG, and EEG

We constructed a unified analysis pipeline (Fig. 1) to compare fMRI, MEG, and EEG using the same THINGS stimulus set (1,854 object categories; Fig. S1a, b). To perform decoding analysis, we first trained modality specific encoding models on layer wise DNN feature representations, with hyperparameters optimized by cross-validation. Among the DNNs considered, for this initial benchmark, we focused on AlexNet (Krizhevsky et al., 2012) as a reference model, because its hierarchical architecture, with progressively more abstract features from shallow to deep layers, provides a natural benchmark for assessing correspondences with the spatial hierarchy observed in fMRI and the temporal hierarchy observed in MEG and EEG (Greene and Hansen, 2018; Wagatsuma et al., 2022; Wen et al., 2018).

Using AlexNet features, the encoding analysis revealed a clear hierarchical correspondence between DNN layers and fMRI responses. Shallow layers (layer1 to layer3) showed the highest prediction accuracies in early visual cortex (V1/V2), whereas deeper layers (layer5, fc6 to fc8) yielded higher accuracies in higher visual areas such as FFA and PPA (Fig. S2). These results replicate the well-established mapping of shallow to deep model features onto early to high-level visual cortex (Cichy et al., 2016a; Güçlü and van Gerven, 2015; Yamins et al., 2014).

In contrast, MEG and EEG exhibited a shared temporal profile characterized by a prominent prediction peak at approximately 100 to 200 ms after stimulus onset followed by a gradual decay (Fig. S3, S4). This temporal window is consistent with prior MEG/EEG studies showing rapid emergence of object information within the first 200 ms (Cichy et al., 2016b, 2014; Gifford et al., 2022). Both modalities showed decreasing prediction accuracy from shallow to deep model layers, consistent with the interpretation that early DNN features best capture the transient feedforward dynamics reflected in early electrophysiological responses. EEG also exhibited a biphasic peak structure (Grootswagers et al., 2022). However, its prediction accuracies were overall lower than those of MEG, and layer distinctions were weaker, reflecting its lower SNR and reduced sensitivity to detailed representational structure.

### Stage-dependent effects of group-level averaging on decoding performance

To systematically assess how the stage of averaging influences decoding performance, we compared five aggregation strategies that operate at increasingly abstract stages of the analysis pipeline. Wave-averaged EEG (wave-avg) averages raw channel × time waveforms across participants, whereas weights-averaged EEG (weight-avg) averages encoding model weight vectors to derive a shared mapping from features to signals. Neither method yielded meaningful improvement, suggesting that averaging raw signals or model weights does not sufficiently suppress idiosyncratic noise. In contrast, sim-avg EEG consistently produced the strongest gains, indicating that transforming responses into a similarity representation before averaging preserves meaningful stimulus relationships while eliminating amplitude variability. We further evaluated two decoding level aggregation methods. Log-likelihood-averaged EEG (ll-avg) initially improved with increasing participant numbers but degraded beyond a certain sample size (Fig. S8), revealing that ll-avg is sensitive to subject-specific variability and unstable when likelihood scales differ markedly across individuals. Rank-averaged EEG (rank-avg), which aggregates only the rank ordering within each likelihood vector, exhibited greater stability but showed no large difference in performance compared to sim-avg EEG. Taken together, these results show that averaging at the level of raw waveforms (wave-avg), encoding weights (weight-avg), or continuous likelihoods (ll-avg) does not reliably enhance EEG decoding performance. Instead, averaging at the level of stimulus relationship structure using sim-avg EEG or rank-avg EEG uniquely leverages the practical advantage of large EEG sample sizes to produce a robust, medium SNR group-level representation (Fig. S9).

In addition to EEG, we examined whether analogous group-level averaging strategies improved decoding performance in fMRI and MEG (Fig. S10). For fMRI, the three decoding level aggregation methods, log-likelihood averaging (ll-avg), similarity averaging (sim-avg), and rank averaging (rank-avg), all outperformed the single-subject baseline and yielded nearly identical performance levels. This indicates that, even in high SNR modalities such as fMRI, transforming responses into stimulus-relationship structure provides a robust and effective means of extracting group-level information. In MEG, where trial-level noise is lower than in EEG but higher than in fMRI, the ordering of aggregation methods reflected their relative information richness, rank-avg

< sim-avg < ll-avg. This pattern is consistent with the interpretation that aggregation based on likelihood retains the most detailed decoding evidence by integrating continuous likelihood values. In contrast, methods based on similarity or rank preserve a progressively coarser relational structure. Taken together, these results show that stimulus relationship averaging is broadly beneficial across modalities. In moderate SNR regimes such as MEG, aggregation methods that preserve richer likelihood information (e.g., ll-avg) yield the strongest gains. It is worth noting that EEG was acquired from 46 participants, in contrast to fMRI and MEG, which involved only 3 and 4 participants, respectively.

### Architecture generalization of encoding patterns

To test generalization of the AlexNet based encoding results, we evaluated additional models and performed encoding analyses for all modalities using features extracted from each architecture. In fMRI, we again found the hierarchical correspondence previously observed with AlexNet: shallow features best predicted early visual cortex, whereas deeper features best predicted higher visual areas, and this pattern was reproduced for all models except MambaOut and InternVL3.5 (Fig. S2). In MEG and EEG, we also observed a common temporal profile in which prediction accuracy peaked around 100 to 200 ms after stimulus onset and then gradually decayed (Fig. S3, S4). For all models except MambaOut and InternVL3.5, prediction accuracy decreased systematically from shallow to deep layers. Although EEG showed lower overall accuracy than MEG and a biphasic temporal pattern, the relative decline in accuracy across layers was qualitatively similar across architectures. Across all modalities, the models showed broadly similar ability to predict brain responses, and none exhibited an extreme advantage. Taken together, these observations suggest that the spatial and temporal relationships between brain responses and DNN features during visual stimulation are not specific to AlexNet. Instead, they may be shared to some extent across different architectures, including CNNs, Transformers, self-supervised models, and vision-language models (Cichy et al., 2016b; Güçlü and van Gerven, 2015; Nakagi et al., 2024; Oota et al., 2022; Yamins et al., 2014).

| Model | Citation | Layers used | Source | Notes |
|---|---|---|---|---|
| AlexNet | Krizhevsky et al., 2012 (NIPS) | conv1–5, fc6–fc8 | torchvision (ThingsVision) | Pretrained on ImageNet-1k |
| ResNet-50 | He et al., 2016 (CVPR) | Layer1.2, layer2.3, layer3.5, layer4.2, avgpool, fc | torchvision (ThingsVision) | Pretrained on ImageNet-1k |
| ViT-B/16 | Dosovitskiy et al., 2021 (ICLR) | Transformer blocks 0, 2, 4, 6, 8, 10 (MLP outputs) + final ln | torchvision (ThingsVision) | Pretrained onImageNet-21k (fine-tuned on 1k) |
| CLIP ViT-B/16 | Radford et al., 2021 (ICML) | Transformer blocks 0, 2, 4, 6, 8, 10 (MLP outputs) + final norm | timm (ThingsVision) | Trained via web-scale image–text contrastive learning (CLIP, OpenAI) |
| DINOv2 ViT-B/14 | Oquab et al., 2023 (CVPR) | Transformer blocks 0, 2, 4, 6, 8, 10 (MLP outputs) + final norm | timm (ThingsVision) | Pretrained via self-supervised learning on LVD-142M |
| MambaOut | Huang et al.,2024 (arXiv) | fc2 outputs from blocks 0.2, 1.3, 2.10, 2.19, 2.29, 3.2 | timm (ThingsVision) | Pretrained onImageNet-21k (fine-tuned on 1k) |
| InternVL3.5-8B | Wang et al., 2025 (arXiv) | hidden states of transformer layers 6,12,18,24,30,36 (last-token pooling) | HuggingFace (OpenGVLab/InternVL3_5-8B-HF) | Three-stage training: CPT → SFT → RL (multimodal) |

**Table S1. Deep neural network models and feature layers used for encoding and decoding analyses.**

This table lists all DNN models used in the study, including convolutional neural networks (CNNs; AlexNet, ResNet-50), vision transformers (ViT-B/16, CLIP ViT-B/16, DINOv2 ViT-B/14), a Mamba-inspired gated CNN architecture (MambaOut), and a large-scale vision-language model (InternVL3.5-8B). For each model, the specific layers from which feature vectors were extracted are shown, along with their implementation source (e.g., torchvision, timm, or HuggingFace) and training details (e.g., ImageNet-1k pretraining, self-supervised learning, or multimodal alignment). These layer selections were chosen to sample early, intermediate, and deep processing stages within each architecture, providing a comparable set of hierarchical features for cross-modality encoding and decoding analyses.

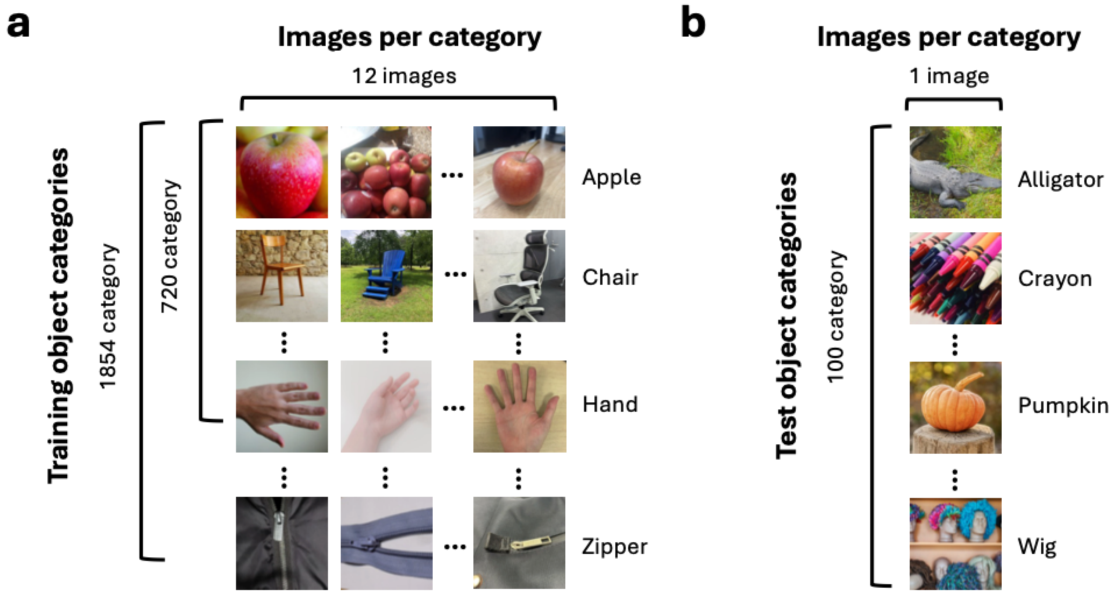


**Fig. S1. Training and test image sets for object-category decoding.**
(a) The training image sets differed across modalities: the fMRI dataset included 720 object categories with 12 images per category (8,640 images total), whereas the MEG and EEG datasets included 1,854 categories with 12 images per category (22,248 images total) (Grootswagers et al., 2022; Hebart et al., 2023, 2019). (b) The test image set consisted of 100 novel images drawn from 100 object categories that were also included in the training set. These images were shared across datasets (fMRI, MEG, and EEG) and were used to compare decoding performance across modalities. For copyright reasons, the images shown here are illustrative substitutes: in panel a, the first image for each category was taken from THINGSplus (CC0) and the remaining images were either CC0 licensed public images or author created images; in panel b, all images were taken from THINGSplus (CC0). These images do not correspond to the actual experimental stimuli (Stoinski et al., 2022).

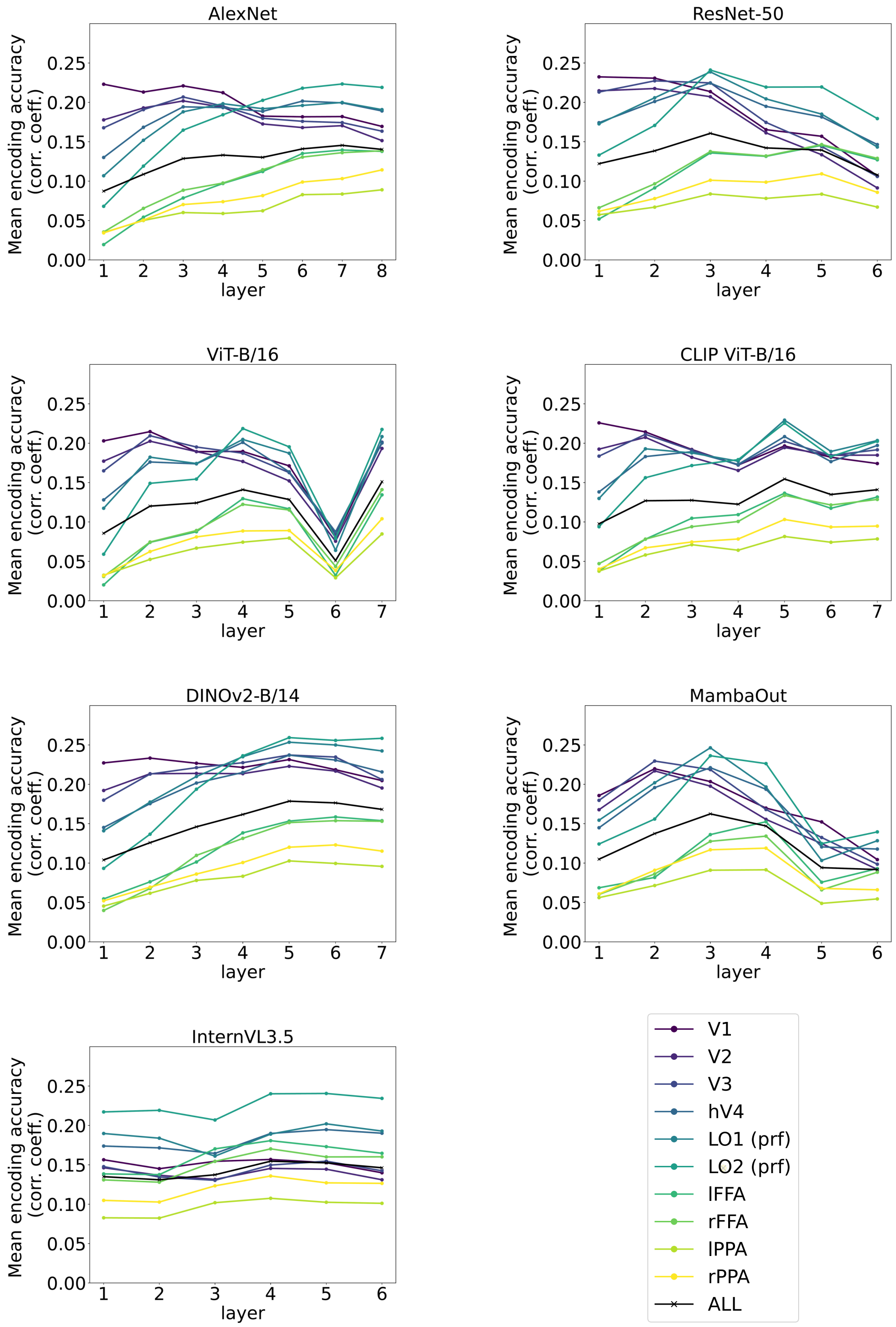


**Fig. S2. Encoding model results for fMRI.**

Voxel-wise encoding accuracy (mean Pearson correlation coefficient) for multiple DNN models across model layers and visual ROIs. Layer-ROI profiles exhibit a clear hierarchical correspondence; early layers align with early visual areas, and deeper layers align with higher visual areas.

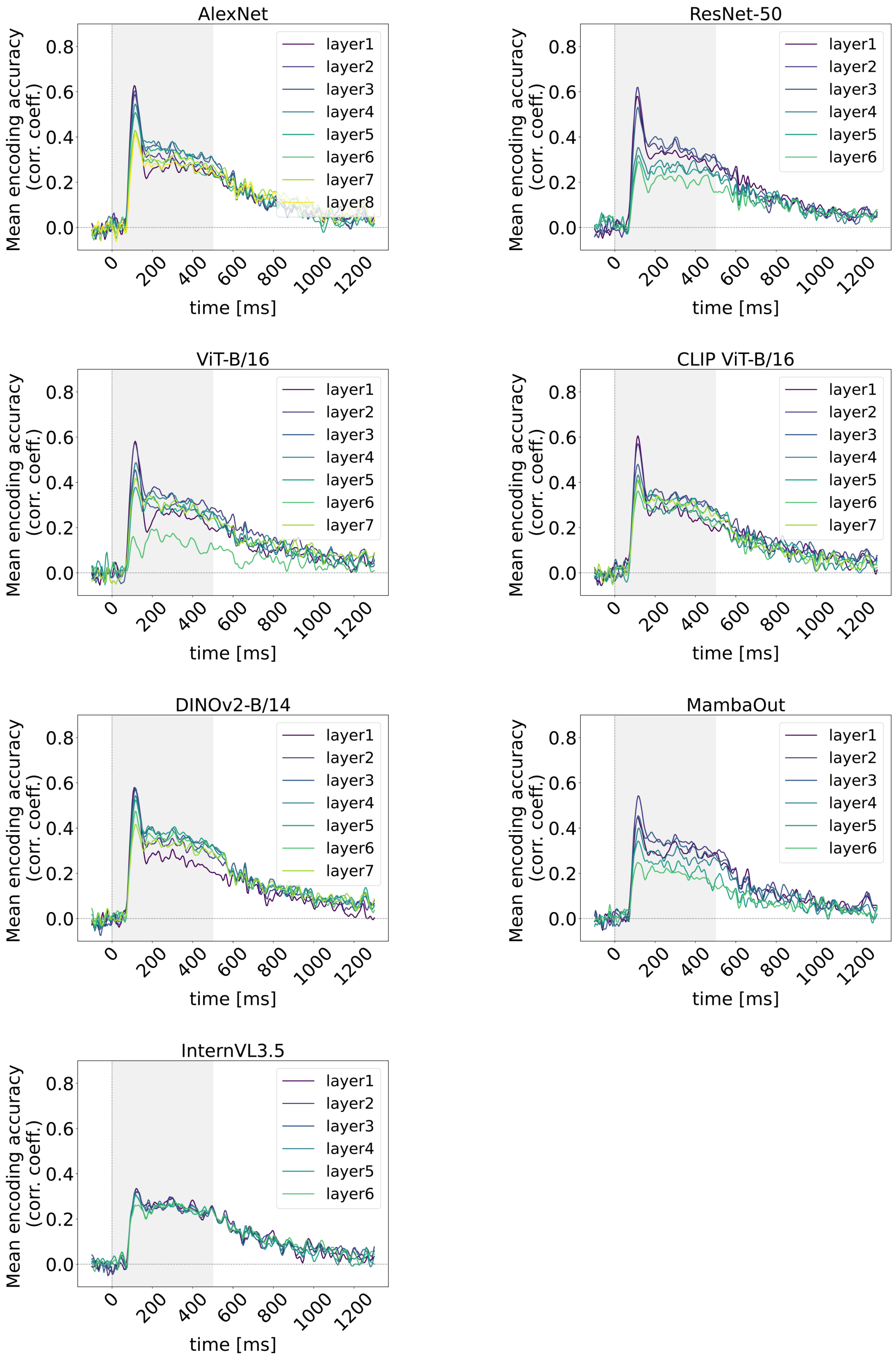


**Fig. S3. Encoding model results for MEG.**

Channel-wise encoding accuracy (mean Pearson corr. coeff.) as a function of time for multiple DNN models and layers. All models exhibit a peak around 100-200 ms followed by gradual decay, demonstrating a consistent temporal profile across models.

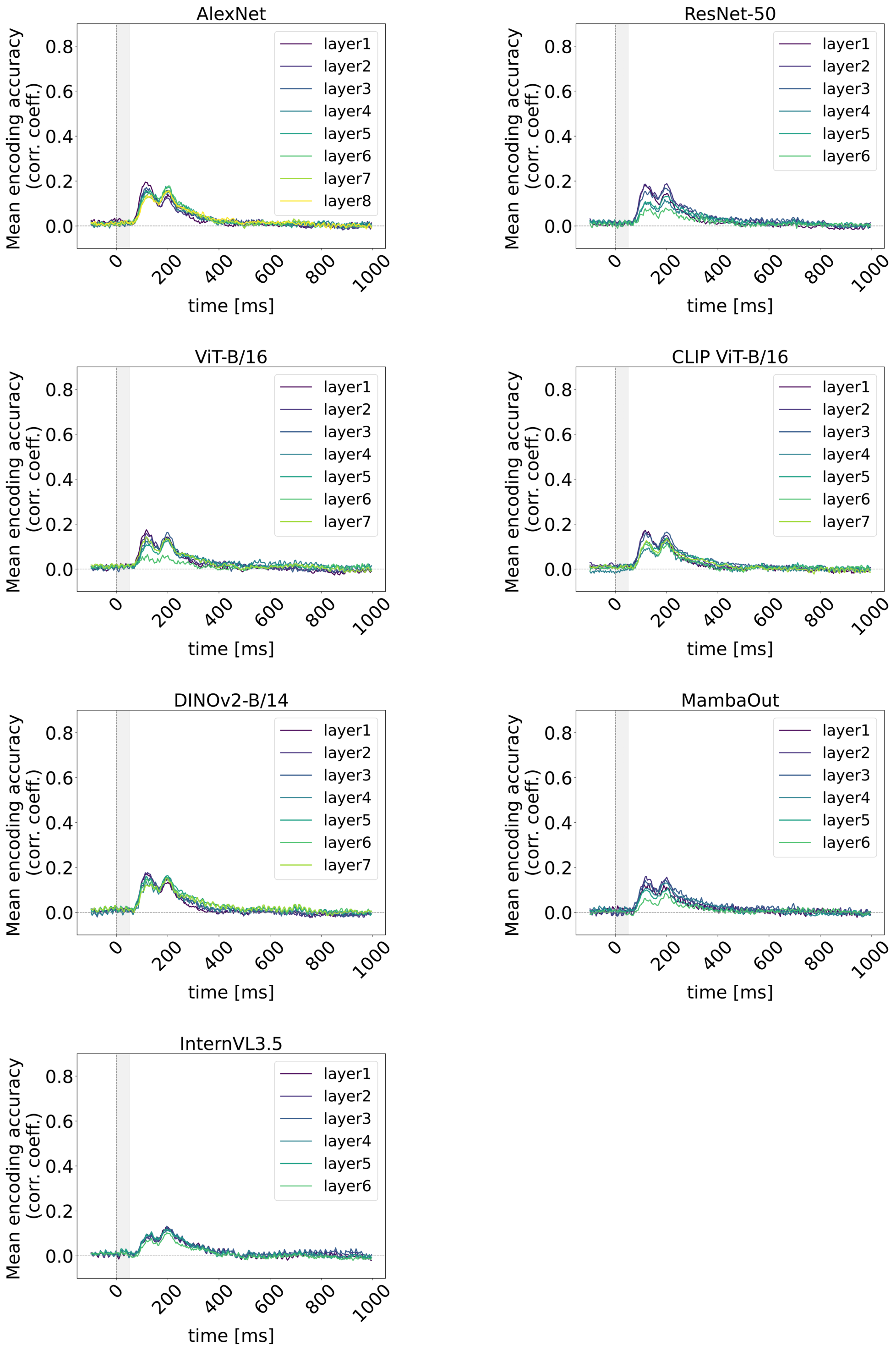


**Fig. S4. Encoding model results for EEG.**

Channel-wise encoding accuracy (mean Pearson corr. coeff.) as a function of time for multiple DNN models and layers. Although absolute correlations were lower than

MEG, the temporal profile and relative decrease from early to deep layers were consistent across models.

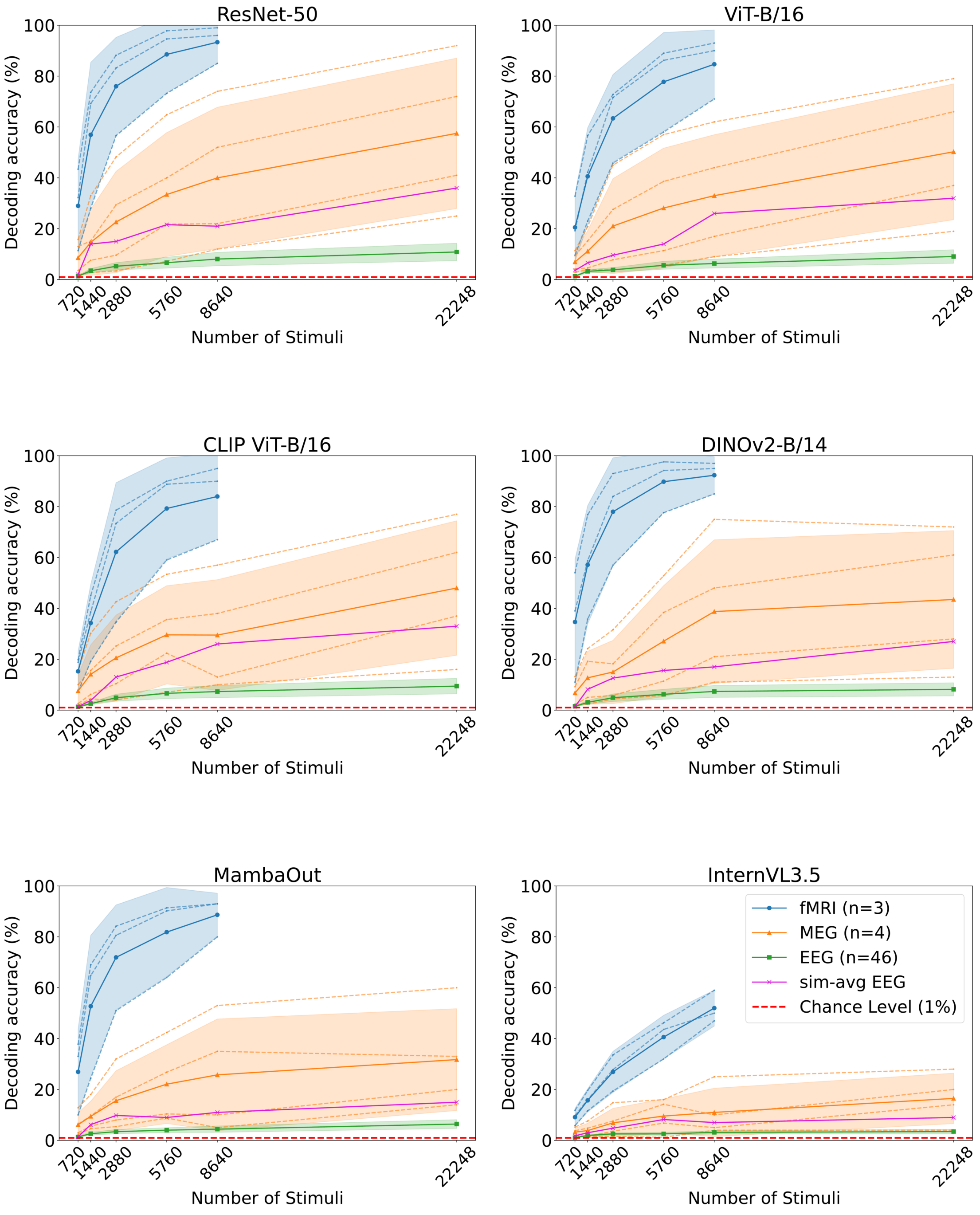


**Fig. S5. Decoding accuracy as a function of stimulus counts across models.**

Decoding accuracy increased with stimulus count for all DNN models, and the relative modality ranking was consistent across models, indicating that the observed modality differences are not driven by model choice. Shaded areas indicate 95% CI, and red dashed lines denote chance level.

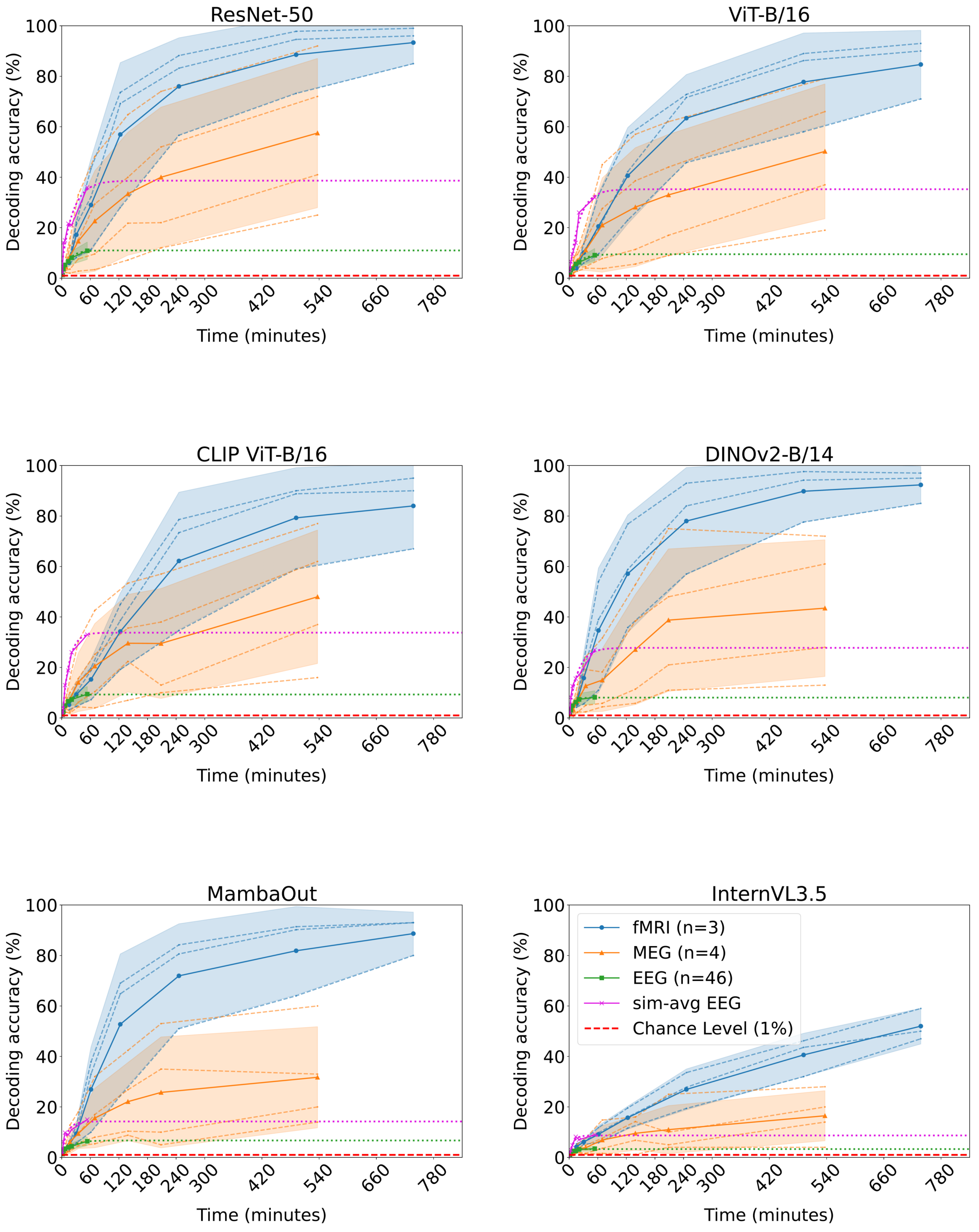


**Fig. S6. Decoding accuracy as a function of measurement time across models.**

Decoding accuracy varied with measurement time across all DNN models, and the efficiency precision trends were consistent across models (sim-avg EEG > MEG > EEG for short durations; fMRI > others for long durations). Shaded areas indicate 95% CI, and red dashed lines denote chance level.

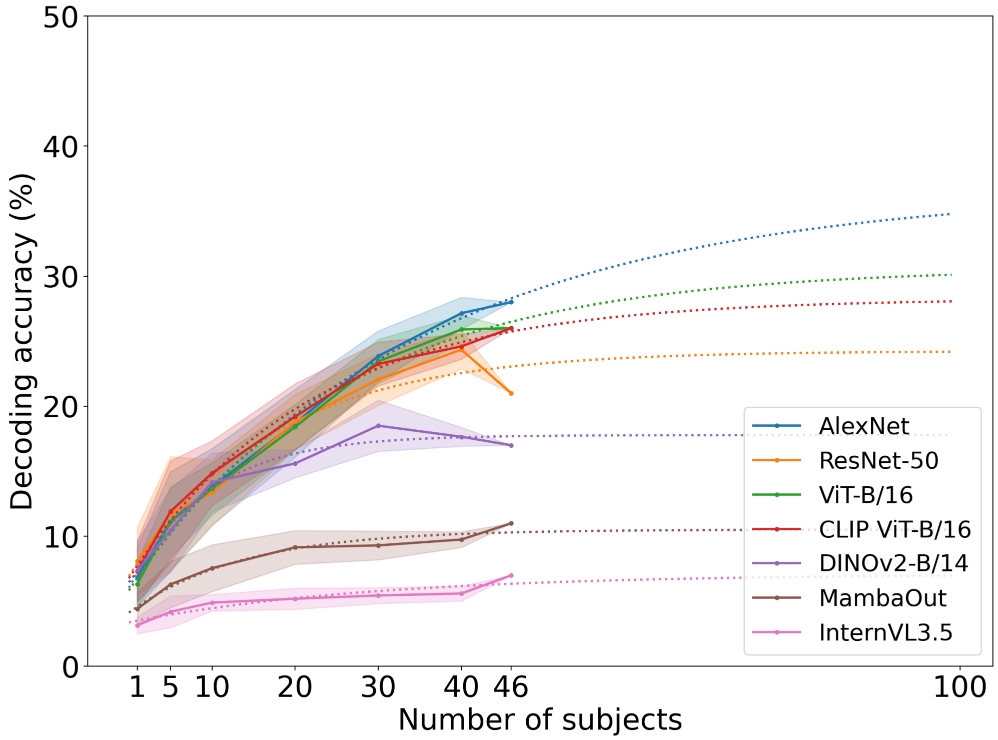


**Fig. S7. Scaling of EEG decoding performance.**

Top-1 accuracy increased with the number of participants included in the group-level analysis (N = 8640). Shaded areas indicate the 95% CI across random subsampling (20 seeds), and red dashed lines denote chance level. A fitted scaling function extrapolated an asymptotic performance of about 36.7% (AlexNet).

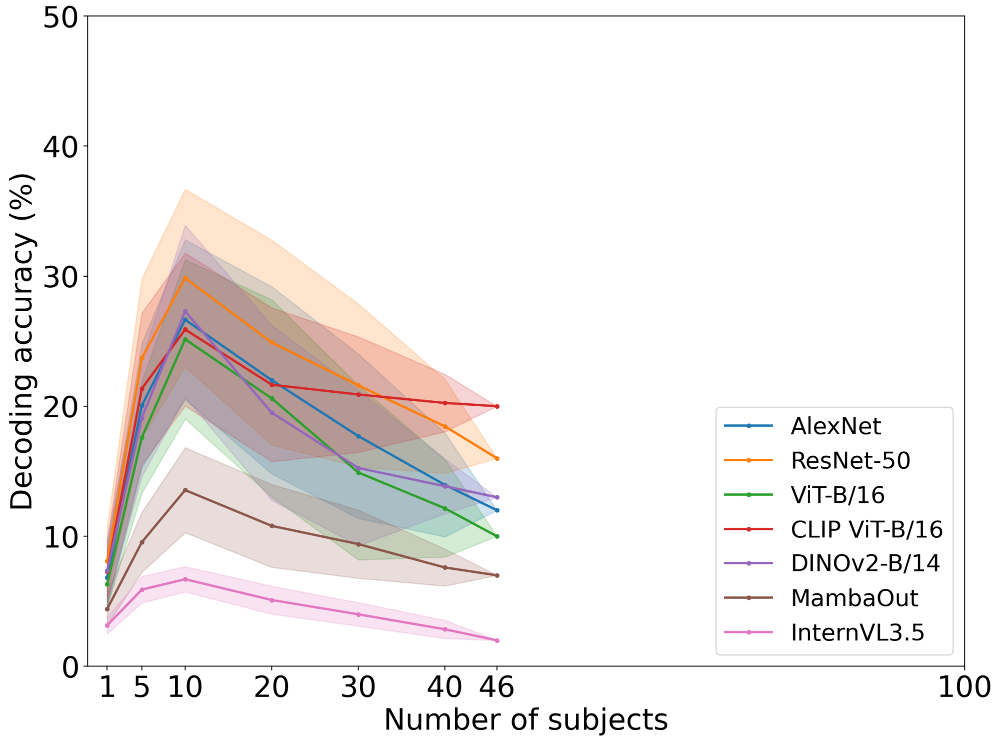


**Fig. S8. Performance degradation in log-likelihood-averaged EEG (ll-avg).**

Although ll-avg EEG initially improved with increasing the number of subjects, performance degraded beyond a certain sample size. This indicates that this averaging method can become unstable as more participants are included (N = 8640). Shaded areas indicate the 95% CI across random subsampling (20 seeds), and red dashed lines denote chance level.

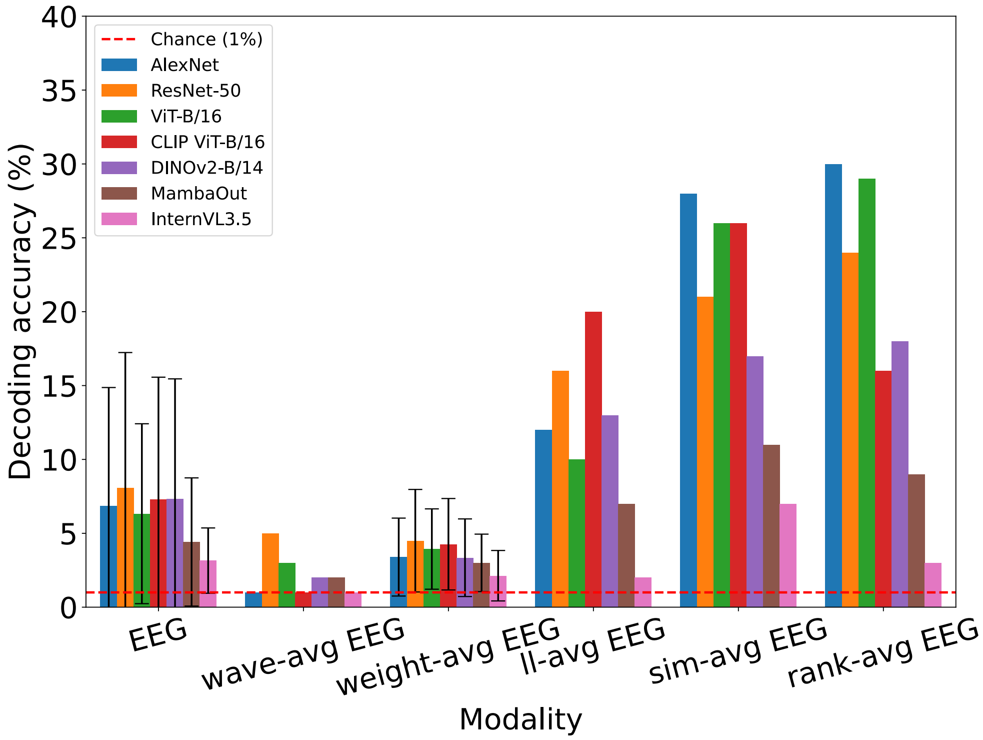


**Fig. S9. Comparison of group-level EEG averaging methods.**

Decoding accuracy for single-subject EEG and five group-level aggregation methods (wave-avg, weight-avg, ll-avg, sim-avg, and rank-avg EEG; N = 8640). sim-avg EEG and rank-avg EEG consistently achieved the highest performance, indicating that appropriate group-level aggregation can substantially improve EEG decoding.

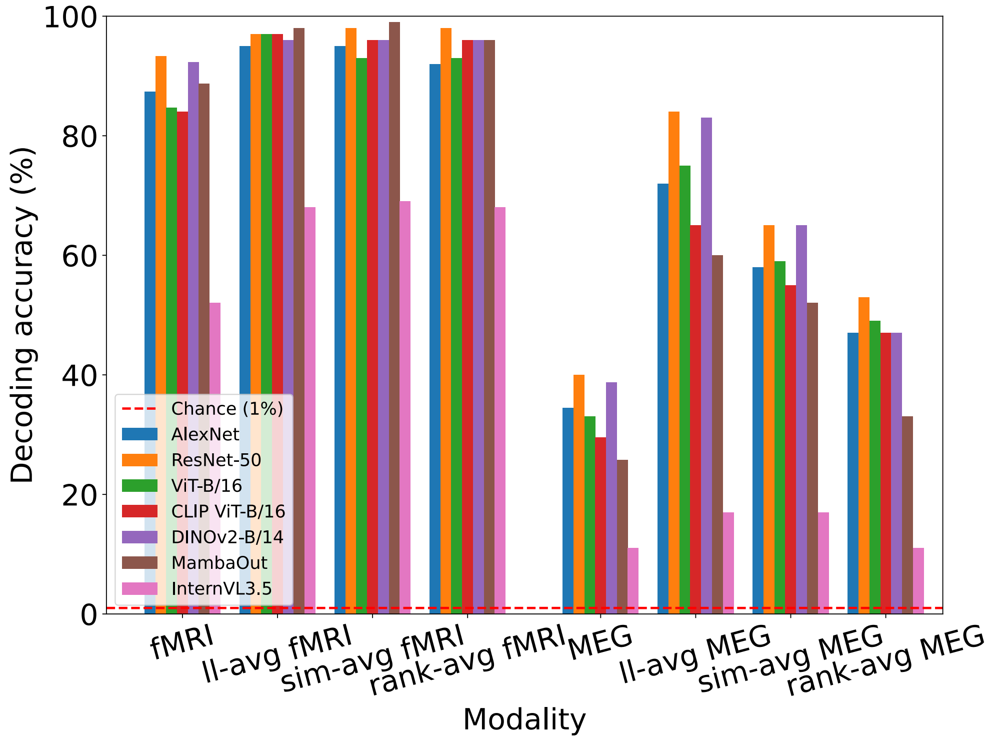


**Fig. S10. Group-level averaging effects in fMRI and MEG.**

Likelihood-based averaging was applied to fMRI and MEG to examine whether the group-level improvement observed in EEG generalizes to other modalities (N = 8640). Both modalities showed modest improvements despite limited sample sizes, suggesting the potential applicability of group-level strategies beyond EEG.
